# Eco-evolutionary feedbacks drive the co-occurrence of restriction-modification systems and antimicrobial resistance genes

**DOI:** 10.1101/2025.06.11.659087

**Authors:** Joseph Westley, Paritosh Bedekar, Elizabeth Pursey, Mark D. Szczelkun, Mario Recker, Stineke van Houte, Edze R. Westra

**Author notes:** Current address: Department of Experimental Medical Science, Faculty of Medicine, Lund University, Lund, Sweden (Elizabeth Pursey). **Corresponding author:** Joseph Westley.

## Abstract

Bacterial pathogens commonly become drug resistant via horizontal acquisition of antimicrobial resistance genes (ARGs), which are often encoded on mobile genetic elements (MGEs). Although bacterial defence systems are typically considered barriers to horizontal gene transfer (HGT), previous studies revealed that bacteria with more restriction-modification (RM) systems (the most abundant bacterial defences) frequently carry more MGEs. It was suggested that this counterintuitive relationship might result from stronger selection for RM systems when exposure to costly MGEs increases. Here, we test this hypothesis using a combination of modelling and bioinformatics analysis of >40,000 bacterial genomes to better understand how eco-evolutionary feedbacks between selection for RM and acquisition of MGEs shape bacterial genome evolution. Our model predicts negative associations between HGT and RM, but only if RM diversity is high. By contrast, at low RM diversity, eco-evolutionary feedbacks drive the emergence of positive associations between HGT and RM. Consistent with these predictions, we identified negative relationships between acquired ARG counts and RM counts across species but positive relationships within individual species. Collectively, our work helps to understand how RM systems shape patterns of HGT of ARGs, which may offer opportunities for targeted surveillance of strains at higher risk of horizontally acquiring novel drug resistance alleles.

**Significance statement:** Previous research shows positive associations between bacterial restriction-modification (RM) systems and mobile genes, despite RM being well-evidenced as a barrier to mobile gene acquisition. Eco-evolutionary feedbacks have been hypothesised to drive this; higher HGT exposure will select for greater RM investment, while still leading to increased mobile gene acquisition. By formalising this hypothesis into a mathematical model and analysing bacterial genomes, we reveal novel insights into how the relationship between RM and mobile antimicrobial resistance genes (ARGs) depends on ecological variables. This study illustrates that to better comprehend how defence systems regulate gene transfer in natural communities, it is essential to consider both their mechanisms and the complex eco-evolutionary dynamics at play.

## Introduction

It is difficult to overstate the urgency of the antimicrobial resistance crisis, with global clinical and agricultural use of antimicrobials on the increase^1,2^, and predictions of 10 million deaths annually from drug-resistant infections for 2050^3^. The recent growth in the prevalence of multi-drug-resistant pathogens is to a large extent driven by horizontal gene transfer (HGT) of antimicrobial resistance genes (ARGs) by mobile genetic elements (MGEs)^4–7^. MGEs are pieces of DNA that specialise in moving between and within genomes, and include plasmids, transposons, integrative and conjugative elements and bacteriophages (phages).

Prokaryotes have evolved a diverse arsenal of defence systems to protect them against infection from MGEs^8^. It remains unclear how important these defence systems are in controlling how ARGs are horizontally transferred. Of the >100 microbial defence systems discovered to date^9^, the most ubiquitous are restriction-modification (RM) systems^10^. Understanding the role RM plays in regulating HGT of ARGs is essential to better understand how multi-drug resistance is acquired, which taxa may be important sources of ARGs, and which taxa may be more likely to acquire them.

RM systems perform two main functions: the methylation of self-DNA at specific target sites (modification), and the degradation of DNA that is unmethylated at these sites (restriction)^11^. Through these two processes, RM systems allow discrimination between self and non-self-DNA and cleavage of the latter, providing the cell with innate immunity against unmethylated MGEs^11–13^. Importantly, if a cell donating an MGE (donor) possesses an RM system that methylates the same site as the RM system of a cell receiving the MGE (recipient), the recipient’s RM system will not cleave the MGE. This is because the donor’s own methyltransferase will methylate the MGE at the target site, protecting the MGE from cleavage by the respective endonuclease, allowing the MGE to escape the recipient’s RM system upon transfer. *De novo* escape can also occur at low frequencies when the recipient possesses an RM system the donor does not, if the MGE is methylated by the recipient’s own methyltransferases before the respective endonucleases can degrade the MGE.

The (imperfect) barrier that RM systems pose to phage and plasmid infection has been demonstrated experimentally^14–27^. Moreover, the role of RM as a barrier to MGE infection is further supported by signatures of restriction avoidance in MGEs. Specifically, the target motifs that RM systems recognise are short (typically 4-8bp for Type II and Type III systems)^28^ but frequencies in many MGEs (especially small plasmids and phages) are often below what would be expected by chance^29–31^ or are in orientations that preclude cleavage^32^. Additionally, anti-restriction systems have been observed in both phages^33–35^ and plasmids^36–38^, and function to neutralise their host’s RM system(s)^11,39^. Furthermore, HGT of ARGs between phylogroups (genetically distinct lineages) in methicillin-resistant *Staphylococcus aureus* (MRSA) has been proposed to be regulated by between-phylogroup variation in RM systems^29^. Similarly, inter-phylogroup rates of homologous recombination in *Neisseria meningitidis* appear to be reduced by RM systems^40^.

Given this extensive evidence that RM systems can pose a barrier to the acquisition of MGEs, intuitively one would predict that lineages associated with more RM systems would carry fewer horizontally acquired genes. However, a bioinformatic study of >800 genomes that investigated the relationship between the presence of RM systems and gene gain via HGT and homologous recombination demonstrated that lineages associated with more RM systems were also associated with *greater* gene gain^41^.

Several potential explanations for these positive associations between RM and MGE content have been put forward, but none of these has been rigorously tested. First, it has been suggested that positive associations may arise as a result of eco-evolutionary feedbacks: as exposure to costly MGEs increases, so will selection for investment into RM systems^41^. A second explanation is that MGEs may carry RM systems (genetic linkage)^42–45^, thus resulting in positive associations. Thirdly, as RM systems can be horizontally acquired^42–45^, it is possible lineages acquired MGEs before they acquired RM, and then these lineages carrying both adaptive MGE-borne genes ^46^ as well as genome defences (RM) may be under strong positive selection.

To identify the potential mechanisms that drive previously reported positive associations between RM and MGEs, we first developed a mathematical model of the eco-evolutionary dynamics of bacterial populations carrying RM systems exposed to MGE infection. This model predicts conditions that drive either positive or negative associations between RM and MGE content, depending on the population-level diversity of RM systems. Next, we tested model predictions through in-depth bioinformatic analyses of >40,000 publicly available genomes of key bacterial pathogens. Due to the greater scale of our dataset compared with previous research^41^, we were able to conduct both between-species and phylogenetically controlled within-species analyses.

Combining these modelling and bioinformatics methods we conclude that eco-evolutionary feedbacks between MGE exposure and investment in genome defence form the most parsimonious explanation for the observed positive relationships between RM and MGE content in bacterial genomes. We propose that this information can be used to identify species networks where HGT is potentially less restricted by RM, which may offer opportunities for targeted intervention strategies to limit the spread of AMR.

## Results and Discussion

### Positive associations between RM and MGEs arise when population-level RM diversity is low

To formalise the hypothesis that eco-evolutionary feedbacks can drive positive associations between RM and MGE content in bacterial genomes, we utilised a modelling approach (see Materials and Methods, Mathematical modelling). We competed three sub-populations with varying levels of investment into RM: zero, low, and high. We allowed the gain of up to five MGEs, with higher investment into RM resulting in a reduced likelihood of MGE acquisition. We varied the rates of HGT as well as the diversity of RM systems in the population, as we predict RM becomes a weaker barrier to HGT when the probability that donors and recipients share the same RM specificity increases^47,48^, thus increasing the probability of gene exchange between lineages.

In simulations where population-level RM diversity was low, bacteria with high RM investment also had a higher mean count of MGEs (Figure 1A), whereas a negative relationship between RM investment and MGE count was observed when the simulated population-level RM diversity was higher (Figure 1B). To ensure robustness of these results, we ran simulations with varying RM investment costs and varying MGE costs. This revealed qualitatively similar outcomes unless RM diversity was low and carrying these systems was more costly than being infected by MGEs (Figure S1).

**Figure 1.**
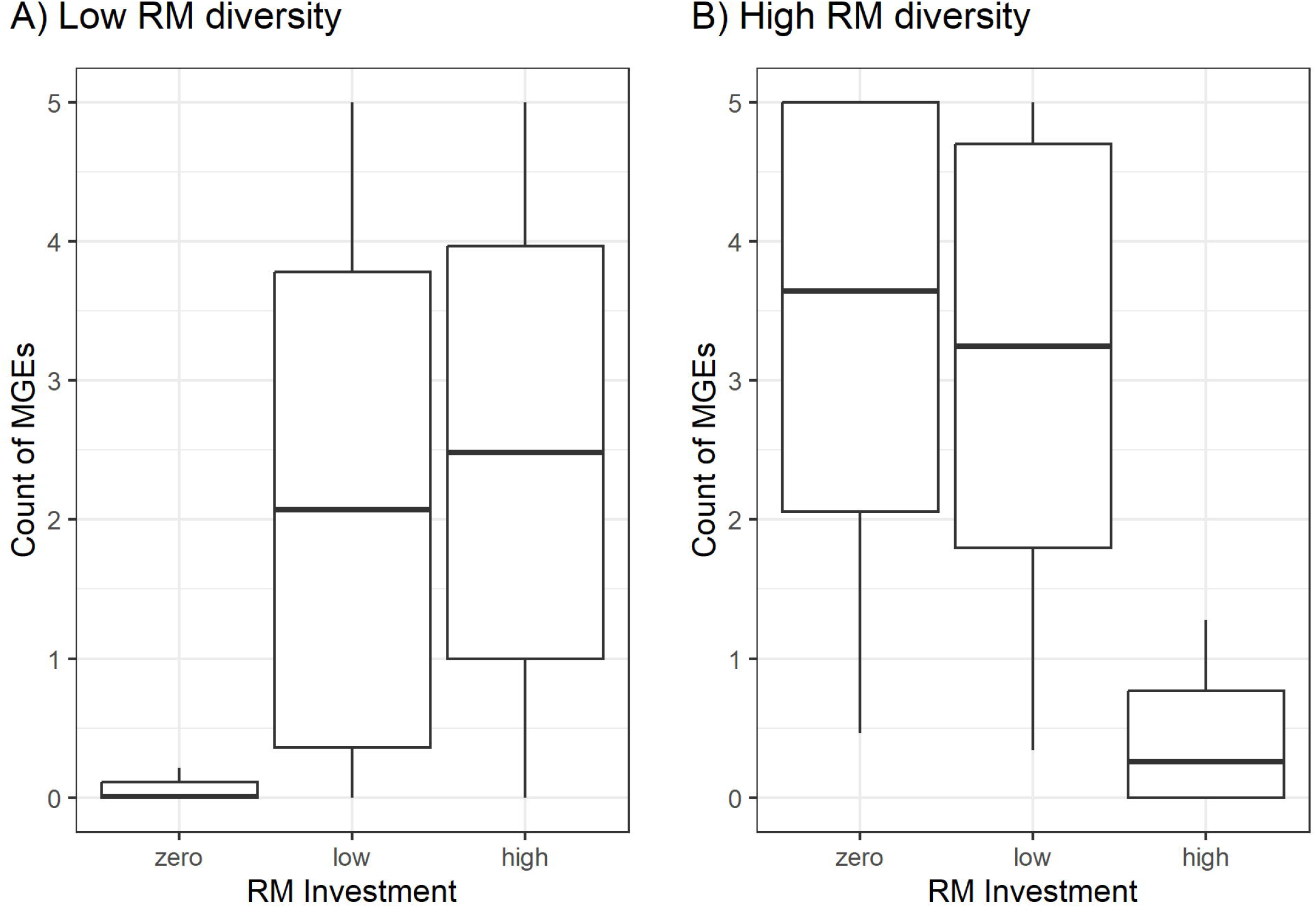
The average count of MGEs carried by populations with different levels of RM investment. Simulations when population-level RM diversity is either A) low or B) high. The central horizontal line indicates the mean, the boxes indicate ± 1 standard deviation (SD) from the mean, the whiskers denote ±2 SD from the mean. Here, data from all simulations across the full range of relative HGT rates are combined.

Collectively, these simulations suggest high exposure to costly MGEs can drive both an increased investment into RM-mediated genome defence as well as an increase in the acquisition of mobile genes, provided that the population-level diversity of RM systems is low. This is likely because RM systems are an imperfect barrier, and once the barrier is crossed, methylated MGEs will be able to infect other bacteria that carry the same RM specificity, but not bacteria with a different RM specificity. In other words, RM diversity impacts the prevalence of routes for gene exchange between lineages with matching RM target sites (as described by Oliveira *et al*^41^), and will therefore impact the directionality of the relationship between MGE acquisition and RM.

### Identification of RM systems and mobile ARGs in a large-scale genomic dataset

To test our model predictions, we utilised bioinformatic analyses to investigate the association between RM systems and horizontally acquired ARGs, leveraging the wealth of publicly available genomic data for bacterial pathogens and established tools for ARG annotation.

We searched a total of 40,181 genomes from 14 bacterial pathogen species from NCBI’s *RefSeq* database, for defence systems and acquired ARGs (Table 1). This 50-fold increase in genomic dataset size, compared with previous analyses of the relationship between RM and mobile genes^41^, allowed us to conduct between-species and phylogenetically controlled within-species analyses.

**Table 1.** Count of genomes analysed, RM systems found, and ARGs found, for each species.

| Species | Genomes | Type I RM | Type II RM | Type III RM | Type IV RM | Total RM | ARGs |
| --- | --- | --- | --- | --- | --- | --- | --- |
| *Pseudomonas aeruginosa* | 4,751 | 1,991 | 1,132 | 370 | 51 | 3,544 | 33,826 |
| *Acinetobacter baumannii* | 3,986 | 574 | 135 | 88 | 1 | 798 | 41,996 |
| *Enterococcus faecium* | 1,947 | 297 | 154 | 94 | 1 | 546 | 22,258 |
| *Streptococcus pyogenes* | 1,753 | 1,687 | 72 | 0 | 0 | 1,759 | 782 |
| *Neisseria gonorrhoeae* | 622 | 498 | 3,253 | 579 | 0 | 4,330 | 97 |
| *Campylobacter jejuni* | 1,478 | 577 | 2,530 | 267 | 0 | 3,374 | 1,956 |
| *Helicobacter pylori* | 1,512 | 1,652 | 5,566 | 1,479 | 0 | 8,697 | 35 |
| *Shigella flexneri* | 415 | 407 | 2 | 0 | 2 | 411 | 3,309 |
| *Shigella sonnei* | 994 | 30 | 982 | 15 | 8 | 1,035 | 6,734 |
| *Clostridioides difficile* | 1,717 | 1,337 | 459 | 4 | 8 | 1,808 | 10,329 |
| *Klebsiella pneumoniae* | 5,184 | 2,826 | 696 | 882 | 186 | 4,590 | 37,306 |
| *Staphylococcus aureus* | 4,669 | 692 | 1,342 | 66 | 3,438 | 5,538 | 27,558 |
| *Mycobacterium tuberculosis* | 5,241 | 0 | 7,875 | 0 | 9,348 | 17,223 | 12,388 |
| *Salmonella enterica* | 5,912 | 4,503 | 773 | 5,835 | 2,581 | 13,692 | 9,804 |
| *Total* | 40,181 | 17,071 | 24,971 | 9,679 | 15,624 | 67,345 | 208,378 |

Across our dataset, we observed systems from over 141 distinct defence system types, as defined by *padloc*, a tool that identifies bacterial defence systems in sequence data. We grouped all functionally similar systems, and of the resulting 41 distinct groups, we found that RM systems were the most abundant type of defence system, with 67,345 complete systems observed (Table 1, Figure S2.1).

Here we further improve upon past work by including all types of RM systems in our analyses, rather than just Type II^41^. The average per genome counts of RM systems varied considerably across species, from 0.20 in *Acinetobacter baumannii* to 6.96 in *Neisseria gonorrhoeae*. For the majority of species, the most abundant RM systems belonged to either Type I or Type II, with Type III and IV systems being less frequent (Table 1), a finding consistent with previous literature^8^. There were however, two outlier species in this regard; over half of all identified Type III systems were observed in *Salmonella enterica*, and over half of all identified Type IV systems were observed in *Mycobacterium tuberculosis* (Table 1).

Across all genomes we also identified 208,378 acquired ARGs. Again, we observed considerable variability in the average counts of ARGs per genome between-species, from 0.02 in *Helicobacter pylori* to 10.54 in *Acinetobacter baumannii* (Figure S2.2).

### Species with more RM systems have fewer acquired ARGs

Firstly, to investigate the relationship between RM systems and acquired ARGs at the broadest scale, we modelled this relationship using data from all 14 bacterial species (see Table 1), using RM system and ARG counts per genome as the explanatory and response variables, respectively. We took a random subset of 400 genomes from each species for this analysis to reduce biases in our data. We found that when we did not control for species identity, there was a strongly negative non-linear relationship observed (Figure 2A, Table S1). However, when species identity was controlled for, this relationship became weakly positive (Figure 2B, Table S1). This suggests that the initial negative relationship observed is driven by between-species differences, i.e., species with more RM systems tend towards having fewer ARGs.

**Figure 2.**
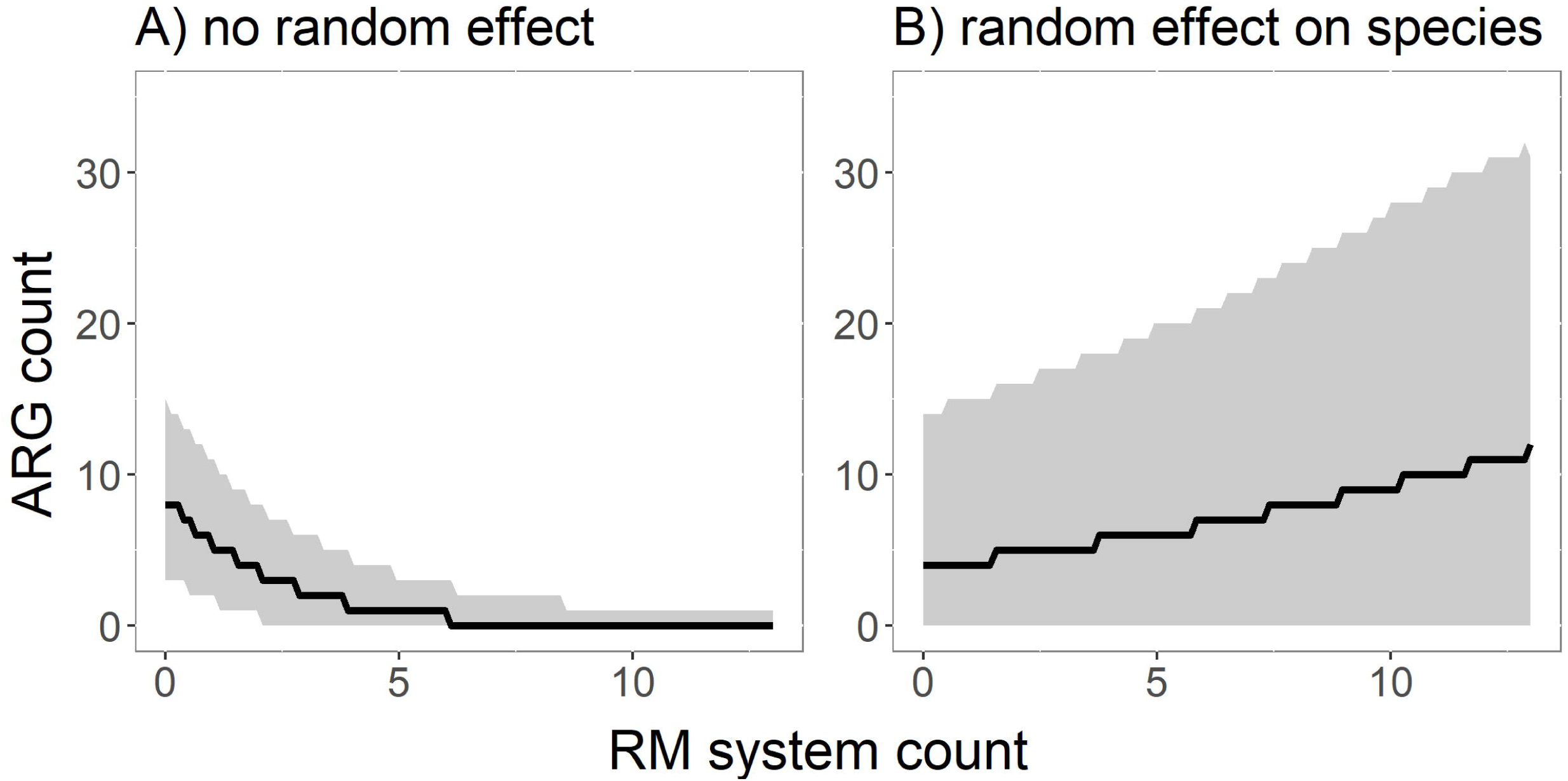
Relationship between the count of RM systems per genome and the count of ARGs per genome as predicted by Bayesian models fit to all taxa data either without (A) or with (B) a random group effect fitted to species identity. Shaded grey area indicates the space within which 95% of predicted draws from the posterior distributions fall. Note, to prevent species with more genomes having a disproportionate influence on the model predictions a dataset comprising random samples of 400 genomes from each species was utilised.

### Within species, genomes with more RM systems have more acquired ARGs

To investigate the relationship between RM systems and acquired ARGs at a finer scale we conducted within-species analyses for five select species from our larger dataset (*Pseudomonas aeruginosa*, *Acinetobacter baumannii*, *Enterococcus faecium*, *Streptococcus pyogenes*, and *Neisseria gonorrhoeae*), controlling for the effect of phylogeny and genome length. These species were chosen as they are taxonomically diverse, all belong to different families, span three phyla, and span the full range of both ARG and RM system counts per genome seen in the full dataset (see Figure S3.1, Figure S3.2, Figure S3.3, Figure S3.4, and Figure S3.5 for RM and ARG distribution across phylogenies).

Consistent with the results of the species identity-controlled global analysis, for four out of five species (*Pseudomonas aeruginosa*, *Acinetobacter baumannii*, *Enterococcus faecium*, and *Streptococcus pyogenes*) the predicted relationships between RM systems and ARGs were positive and had 95% credibility intervals that did not span 0 (Figure 3, Table S1), meaning the models have a high degree of confidence the true relationship is positive. The effect size of RM on ARG count was much larger for *S. pyogenes*, than for the other three species. For *Neisseria gonorrhoeae*, the predicted relationship trended towards being positive, but the 95% credibility interval did span 0 (Figure 3, Table S1). It is notable that we had much broader posterior distributions for our predictions for *Neisseria gonorrhoeae*, suggesting uncertainty. This could be due to reduced statistical power, owing to it being the species with the fewest genomes and fewest observed ARGs per genome in our dataset.

**Figure 3.**
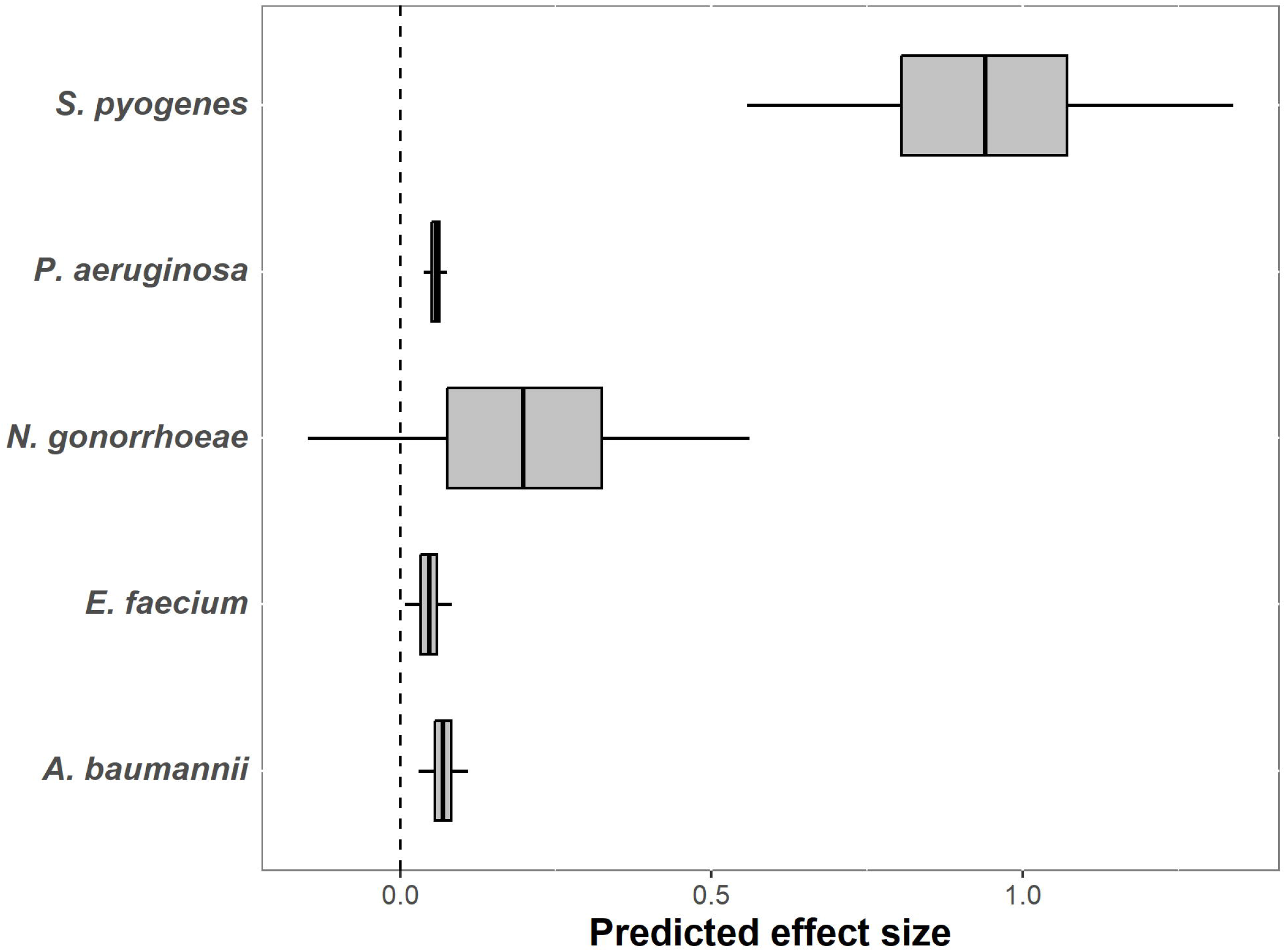
Draws from posterior distributions of the predicted effect of RM system count on ARG count per genome, after controlling for phylogeny and genome length. The boxes denote the interquartile ranges, and the whiskers denote the 95% credibility intervals of the predicted effect sizes.

### Between– and within-species relationships between RM and MGEs differ due to differences in diversity of RM

We observed that between species there are negative relationships between the ARG count and the number of RM systems, while within species, we consistently observe positive relationships. Our mathematical modelling predicts that these results may emerge as a consequence of differences in the RM diversity levels across these scales. We therefore hypothesised that as RM system richness between species will necessarily be higher than within species, RM systems will pose a more effective barrier to HGT of ARGs between heterospecifics than between conspecifics.

To test this, we first analysed the levels of RM system diversity between and within species. We show that all individual species have significantly lower RM system Shannon diversity indices than the entire dataset combined (Table S2). Next, we calculated for every RM system observed in a species the probability that it would restrict intra-versus interspecific HGT events, based on the sequence similarity between RM systems in each genome. Within species, for the five species we conducted intraspecific bioinformatic analyses for, the average probability that a recipient cell will have an RM system that a donor cell does not is 0.69 ± 0.31. Between species, for all fourteen species in the analyses, the average probability that a recipient cell will have an RM system that a donor cell does not is 0.96 ± 0.01. Hence, this suggests that on average RM will more frequently function as a barrier to HGT between species than within species (see Table S3 for all within-species probabilities and all pairwise between-species probabilities), resulting in positive and negative associations between ARGs and RM system counts within and between species, respectively, as predicted by our mathematical model.

### ARGs and RM rarely co-localise on the same MGE

An alternative hypothesis for the positive associations we observed between ARGs and RM within species is that specific RM systems and specific ARGs could be co-localized on the same MGE and acquired together by the same lineage. To investigate the prevalence of this, we identified specific ARG and specific RM system pairs that co-occurred at much higher rates than would be expected by chance. 147 unique pairs of RM systems and ARGs were observed to significantly co-occur (Bonferroni-corrected *P* values <.05, Table S5). The percentage of genomes carrying a linked pair varied notably across species (*Pseudomonas aeruginosa* = 40.18%, *Acinetobacter baumannii* = 12.32%, *Enterococcus faecium* = 6.27%, *Streptococcus pyogenes* = 1.03%, and *Neisseria gonorrhoeae* 10.13%). To test whether co-occurrence resulted from co-localization, we evaluated the genetic distance between co-occurring pairs, the frequency at which pairs were observed on the same contig, and genomic context of the pair (see Materials and Methods: ‘Identification of co-localisation of specific RM systems and specific ARGs’). From this we identified only one instance where RM-ARG co-occurrence is explained by origin on the same MGE. In this case, the respective Type II RM system and two macrolide resistance genes were co-localised on the previously described *S. pyogenes* conjugative prophage Φ1207.3^49–51^ (Figure S4). This explains why the positive effect size of the relationship between RM and ARG count per genome is much larger for *S. pyogenes* than for the other species (Figure 3). There is a clinical significance to this co-localisation as Type II RM systems can function similarly to toxin-antitoxin (TA) systems^52,53^. As other TA systems have been shown to stabilise ARG carrying MGEs^54^, MGEs that carry a Type II RM system as well as ARGs will be under stronger selection for maintenance than MGEs lacking an RM system. Notwithstanding this example of co-localization, we conclude there is no evidence that RM and ARGs co-localising on the same MGE is driving the positive associations we observed in our within-species analyses. However, it is notable that despite co-localisation of specific RM systems and ARGs being very rare, significant co-occurrence is much more common. This is consistent with the idea that there are channels of increased gene exchange between lineages with the same RM systems. If a novel ARG is horizontally acquired by a lineage with a certain RM system, then it will be more readily transferred to other lineages with this RM system, causing the novel ARG and RM system to co-occur significantly more often than would be expected by chance.

### RM systems are positively associated with ARGs, even when acquired first

We also hypothesised that, as RM systems are themselves often horizontally transferred^42–45^, the order of acquisition of RM systems and ARGs could impact their relationship. For example, if a lineage acquired an RM system more recently than an ARG, the RM system would have no opportunity to pose a barrier to the ARG’s acquisition. To ensure the conclusions we drew from our within-species analyses were not confounded by this, we calculated the ‘trait depth’, a measure of the evolutionary time since genetic material was acquired, for each RM system and each ARG in our dataset (see ‘S2.2 Trait depth filtered modelling’). We then filtered our dataset to include only RM systems that fell within the upper tertile (33^rd^ percentile) for trait depth (the earlier acquired RM systems), and ARGs that fell within the lower tertile for trait depth (more recently acquired ARGs). We then conducted within-species analyses of the relationship between RM systems and ARGs using this trait depth filtered dataset.

For *S. pyogenes*, the relationship between count of RM systems and acquired ARGs per genome, which was positive when utilising all the data, became a null relationship when the model was trained on the trait depth filtered dataset (Figure S5). This is likely because here our filtering removed the two ARGs and RM system associated with the conjugative prophage Φ1207.3. However, for the other four species, the directionality of the ARG∼RM relationship remained unchanged. From this we conclude that our finding that ARGs are positively associated with RM systems within species is robust.

Interestingly, we found that for four out of five species, RM systems in general had a higher trait depth than ARGs, indicating that they evolved or were acquired more distantly in evolutionary time (Figure S6). This further reinforces that even when we do not filter our dataset by trait depth, in any given lineage, RM systems are likely to have been present prior to the acquisition of the respective lineage’s ARGs. This disparity in trait depth between RM and ARGs could be explained by selection for antimicrobial resistance increasing dramatically over the last century with the advent of the clinical use of antimicrobials^55^, while this is likely not the case for selection for defence systems.

### Conclusion and outlook

Here, we combine mathematical modelling and bioinformatic analyses to demonstrate that variation in HGT exposure between lineages can lead to positive associations between mobile genes and RM systems, across a range of realistic parameter values. Additionally, we show that increasing population-level RM diversity increases its efficacy as a barrier to gene flux and thus causes the relationship between RM investment and MGE acquisition to become negative. We also demonstrate that the positive associations we observe within species are not confounded by RM systems and ARGs co-localising on the same MGE, or by lineages acquiring ARGs prior to RM systems.

In this study, we used a broad range of taxonomically and physiologically distinct species that occupy diverse ecological niches, and that are exposed to different antibiotic treatments in the clinic.

Because of this, there is minimal overlap in their ARG content and little evidence of HGT between these species, even in those instances where RM systems are not predicted to form a barrier (Table S3, S4 and Figure S7). Metagenomics studies will be crucial to test how patterns of HGT between co-occurring species in a microbial community are mediated by the RM content of the individual species.

Inevitably, exchange of MGEs will not only depend on bacterial immune systems but also on MGE strategies to overcome bacterial defences^56,57^. Plasmids have been shown to carry many anti-defence systems in the leading region that enters the cell first during conjugation, allowing rapid gene expression and neutralisation of host defences^58^. Of these anti-defences, anti-restriction proteins and protective methyltransferases are some of the most abundant^58^, highlighting the importance of RM systems in the arms race between bacteria and MGEs. Furthermore, experimental study has shown the anti-restriction proteins and methyltransferases of conjugative plasmids allows them to overcome host RM systems^27^ (additionally, the leading region of conjugative plasmids has recently been shown to be depleted in RM target sites^59^). Although our mathematical models did not include anti-restriction, we can speculate that if hosts increase RM system investment in response to rising MGE exposure, MGEs will, in turn, evolve stronger anti-restriction mechanisms. Such an evolutionary arms race could mediate the efficacy of RM systems and contribute further to positive associations between the presence of RM systems and mobile genes.

Here we illustrate that ecological and evolutionary dynamics can lead to counter-intuitive patterns. We note that previous research has shown that RM systems can also promote prophage acquisition, through allowing phage exposed bacterial populations to reach densities that favour phages entering the lysogenic life cycle^60^. Our findings here add to a growing body of evidence that suggests the interplay between defence systems and MGEs is governed not just by molecular mechanisms, but also by the broader ecological context in which these interactions unfold.

In the present study we specifically focused on interactions between RM systems and MGEs, as RM systems are highly ubiquitous, are known to pose a barrier to several forms of HGT and types of MGEs^14–27^. However, the rapid expansion of the known microbial defence repertoire over recent years^9,61^ raises many questions that future research into defence and MGE evolutionary ecology could aim to answer. For instance, how might other defence mechanisms contribute to eco-evolutionary dynamics? An abortive infection system that arrests growth upon infection^62^ could have different implications for evolutionary trajectories compared with systems that degrade MGEs, like RM or CRISPR-Cas. Moreover, how might the specificity of defences impact responses to diverse MGE infections? We hypothesise that the fitness benefit of a broader range defence system, like RM, will depend on exposure to a more diverse range of MGEs, that may span both positive and negative fitness effects. Finally, as single genomes frequently carry several defence systems ^63^, it is necessary to consider how defences interact^64^, from additive effects^65,66^, to synergistic^67–71^ and even antagonistic interactions^72^, to shape the co-evolution of bacteria and MGEs.

There is at present considerable interest in developing a better understanding of how bacterial defence systems interact with and regulate the acquisition of mobile genetic elements. Here we demonstrate that although RM systems have been shown to be barriers to HGT in more simplistic experiments, in complex microbial communities the relationship between RM systems and gene acquisition is more complex. Our work highlights that if we wish to deepen our understanding of how horizontal transfer of genetic material impacts genomic evolution outside of the laboratory, and how defence systems can impact this, we must synthesise knowledge of the mechanistic nature of these processes with an understanding of the ecological and evolutionary dynamics at play.

## Materials and Methods

### Mathematical modelling

To better understand the ecological conditions under which selection could favour positive associations between investment into defence systems and horizontally acquired genes, we built a mathematical model to simulate population dynamics. We considered sub-populations with three levels of investment into RM systems – zero, low, and high. High investment into RM results in a high reduction in likelihood of acquiring an MGE, low investment into RM results in a lower reduction in likelihood of acquiring an MGE, and zero investment into RM results in no reduction in likelihood of acquiring an MGE. Individuals could acquire and lose up to 5 identical MGEs. The rate of acquisition of MGEs was also dependent on the number of recipients (susceptible individuals), the number of donors (individuals with MGEs), the relative rate of HGT, and a contact factor.

The loss of MGEs was given by the product of a basal MGE clearance rate and the number of MGEs carried by the individual. The acquisition and loss of RM systems was not considered. Bacteria had a birth rate ‘β’ and a death rate ‘φ’. Populations with MGEs suffered a reduced birth rate given by (1 – c_mge_)*^i^*where c_mge_ was the cost of one MGE and ‘*i*’ was the number of MGEs present. Populations with an RM system suffered a reduced reproductive rate, given by (1 – c_n_) where c_n_ was the cost of the RM investment strategy. As RM systems do not pose a barrier to MGEs that come from donors with the same RM system, because the MGE will have the protective methylation signature, we partitioned HGT exposure experienced by populations with RM systems into two categories, one from unmethylated MGEs that occurred at a reduced rate, and one from methylated MGEs that occurred without any reduction. The equations governing the population dynamics are given in the Supplementary Information (see S2.1 Mathematical Modelling).

We assumed an initial condition of one individual of each sub-population. With three RM system strategies and six possible MGE counts (from zero to five) we considered eighteen individuals in our initial condition. We simulated the growth of this system until it reached a steady state under different ecological conditions – the cost of the MGEs, the cost of RM investment, and the relative rate of HGT. The cost of MGEs varied between a range of values consistent with previous experimental research and modelling of plasmid costs^73–75^. The costs of the high investment RM system were informed by previous experimental work demonstrating that RM system costs are generally undetectable to low, but can be up to a 13% reduction in birth rate^76^. As the low investment RM system was defined as 64 times less effective at reducing HGT than the high investment RM system, its cost was scaled to always be 64 times less than the respective high investment RM system’s cost. The relative rate of HGT varied between 0 and 1. For each simulation, once it had reached a steady state, we measured the relative abundancies of every sub-population.

We used MATLAB R2023a^77^ for our simulations. We also simulated the acquisition of MGEs under increased population level diversity of RM systems by manipulating the probability an MGE would be methylated with correct methylation signature to confer protection during HGT events.

To ensure our conclusions were not dependent on specific starting parameters, we ran all simulations across a range of MGE and RM costs. For each combination of fixed MGE and RM investment costs, we pooled the relative abundances of all sub-populations from all simulations across the range of relative rates of HGT simulated. See ‘S2.1 Mathematical modelling’ for specific equations and Table S6 for parameter values.

### Bioinformatics methods

#### Study species

As a key motivation for this research is to understand how bacterial defence systems may influence the HGT of ARGs, 14 study taxa (Table 1) were selected based on their listing as either of urgent or serious concern in the CDC’s 2019 Antibiotic Resistance Threats Report^78^. For each species, all complete genomes present within the National Centre for Biotechnology Information’s (NCBI) *RefSeq* database^79^ were downloaded using the command line tool *ncbi-genome-download*^80^ (genomes downloaded August 2022). For our initial global ‘between-species analysis’, all genomes from all 14 species were used (Table 1), but for our within-species phylogenetically controlled analyses we selected five of these species. These species were chosen as they are taxonomically diverse and span the full range of both ARG and RM system counts per genome seen in the full dataset. We also excluded species that lacked heterogeneity in their RM system counts (for example, *Shigella sonnei* and *Streptococcus flexneri* genomes were highly uniform in their RM system content and thus statistical power would be limited).

#### Detection of RM systems

All bioinformatic analyses were conducted using the University of Exeter’s Cornwall based High Performance Computing Cluster, known as Athena. The package PADLOC (Prokaryotic Antiviral Defence LOCator)^81^ was used to search all bacterial genomes for defence system genes using Hidden Markov Models (HMMs). The database used was the most up-to-date version of PADLOC-DB available at the time the research was conducted (August 2022), and all RM system gene HMMs used in this study were authored by L. J. Payne^81^. All genes classified in the PADLOC output as belonging to either a Type I, Type II, Type III, or Type IV RM system were then filtered so only hits where the fraction of the target sequence aligning to the HMM, and the fraction of the HMM aligning to the target sequence were both above 0.9, were retained. These values were chosen after adjusting them and conducting multiple sequence alignments in *Geneious Prime® 2023.0.1*^82^, using the Geneious alignment tool, with random samples of ∼20 genes matching the same HMM, and observing that >95% of them would align, suggesting that they were homologues.

#### Detection of acquired ARGs

Methods for the detection of acquired ARGs were informed by and adapted from the scripts used in Pursey *et al*^83,84^. Specifically, the tool *ABRicate*^85^ was used to search genomes for acquired antibiotic resistance genes, using the NCBI’s *AMRFinderPlus* database^86^.

#### Within-species phylogeny construction

In order to control for phylogenetic relatedness of genomes, rooted phylogenetic trees were created for each species. First, pairwise distance estimation was conducted for using the function *dist* from the package *mash* ^87^, using a k-mer size of 21. Unrooted phylogenetic trees were created from the resulting pairwise distance matrices using the *nj* (neighbour joining) function of the R package *ape*^88^. These trees were then rooted to an outgroup’s *RefSeq* representative genome, using *ape::root*.

Outgroups as close to the ingroup as possible were preferentially selected. Finally, chronograms were created from the rooted trees using *ape::chronos*, the method ‘relaxed’, and a lambda value of 1.

Phylogenetic trees were plotted against presence/absence heatmaps of ARGs and RM systems using the tool *ggtree*^89^ (see supplementary Figures S3.1-S3.5).

#### Calculating diversity indices and probability recipients have RM systems that donors do not

To test if RM diversity within a species was significantly different to the RM diversity between all species, we performed a Hutcheson t-test^90^ on the respective Shannon diversity indices^91^ using the function *Hutcheson_t_test* of the R package *ecolTest*^92^.

We also calculated the probability that a HGT event between two genomes of the same species, and two genomes of different species, will be able to be impeded by the recipient genome having an RM system that the donor genome does not.

For all RM systems observed in a species, we calculated the probability that the RM system will not be able to pose a barrier to HGT between conspecifics by adding together the probabilities that both donor and recipient will have the RM system, neither will have the RM system, and only the donor will have the RM system. We can then find the mean of this value for all RM systems in the species, then minus it from 1 to give the average probability that intraspecific HGT will be impeded by RM. We then did this for all pairwise combinations of different species to get the probabilities that RM will impede HGT between species. Note, this was done twice for each pairwise combination, once where species A is the recipient and B is the donor, and once where species A is the donor and B is the recipient.

#### Identification of co-occurrence of specific RM systems and specific ARGs

To identify instances where a specific RM system and a specific ARG co-occurred more often than would be predicted by chance, the tool Coinfinder^93^ was used, in conjunction with the phylogenetic trees. As all possible RM-ARG pairs were tested for co-occurrence, Bonferroni correction was applied to alpha values used to determine significant co-occurrences, adjusting for n hypotheses where n = ARG richness * RM system richness. This makes our estimations of co-occurrence highly conservative. The default Coinfinder setting of excluding rare elements (ARGs or RM systems) was not utilised.

#### Identification of co-localisation of specific RM systems and specific ARGs

To determine if co-occurring pairs were also co-localised, and thus likely to originate from the same HGT event, we calculated the genetic distance between the RM systems and the ARGs, as well as the frequency at which they were observed on the same contig. To identify pairs that were likely candidates for originating on the same MGE, we conservatively filtered the list of co-occurring pairs to include only those where the mean proximity was <100kb, reducing our 146 pairs to 7 (of these 7 only 3 unique RM systems were observed). A random sample of genomes containing each of these 7 pairs were then selected for further investigation to determine if the contigs in question were potentially extra-chromosomal, or if the regions of DNA containing both RM system and ARG were potentially MGE in origin.

In *Streptococcus pyogenes* there was one instance of an RM system co-localising with two macrolide resistance genes and, based on inspection of the adjacent annotations, this region was suspected of being prophage in origin. To confirm this, the prophage search tool VirSorter2^94^ was run on all genomes possessing the co-localising genes (test), as well as 20 random genomes without these genes (control). All prophage sequences identified from the test genomes were mapped to the largest prophage sequence that spanned the RM system-macrolide resistance gene containing region using Geneious’ built in ‘map to reference’ tool, using the highest sensitivity settings, and iteratively mapping back to the assembly up to 25 times. To confirm this prophage was not present in the control genomes, the final contig produced from this ‘map to reference’ function was then used as a reference to map all suspected prophages from the control genomes.

Two other RM systems were also observed <100kb from an ARG that they were significantly found to co-occur with, but in both instances, there was very limited evidence this was due to origin on same MGE. In *P. aeruginosa* a Type I RM system that was often located next to integron and integrative conjugative element associated genes, and in close proximity to several ARGs. However, due to the high variability in the local sequence in this example and known function of integrons to facilitate incorporation of genetic material into the chromosome^95^, it is probable that here co-localisation is due to this being a ‘hotspot’ of acquired genes, rather than the RM system and ARGs being originally co-localised on the same MGE. In *Acinetobacter baumanii* a Type I system was also observed ∼38kb from a metallo-beta-lactamase, but this co-localisation was only observed 5 times, despite the RM system and ARG being observed 237, and 229 times respectively, and only in the same genome 57 times.

#### Bayesian mixed-effect modelling

All statistical analyses were conducted in R v. 4.3.1^96^, on the University of Exeter’s RStudio pro server. The *tidyverse* packages^97^ were used for data handling. To create figures the R packages *ggplot2*^97^, *ggpubr*^98^, *cowplot*^99^, *RColorBrewer*^100^, *gridExtra*^101^, and *patchwork*^102^ were used. To create tables *flextable*^103^ and *ztable*^104^ were used.

The R package *brms* was used to conduct all statistical analyses^105^, using phylogenetically controlled Bayesian Poisson generalized linear models. All models utilised count of acquired ARGs as the response variable and count of RM systems as the explanatory variable. As we predicted genome length to be positively associated with both ARGs and defence systems, we controlled for it by including it as an additional fixed effect in our models. As *brms* requires a covariance matrix for phylogenetic control, the trees were converted to such using *ape::vcv*^88^. All models were run for 2000 iterations. The default prior intercept used was a uniform distribution with a lower bound of 0 and an upper bound of the highest ARG count per genome from the respective dataset, this is uninformative yet confined to the bounds of what is possible. For *Streptococcus pyogenes*, *Neisseria gonorrhoeae*, and the ‘all taxa’ models, a slightly more informative prior was needed owing to divergent transitions, and a normal distribution with a mean of the actual ARG mean, a standard deviation (SD) of the real ARG SD, and the aforementioned bounds was used. The slope prior was a gaussian distribution with a mean of 0 and a standard deviation of 1. For the all-taxa analysis there was no phylogenetic control, but a random group level effect on species was used. Samples from the modelled posterior distributions were drawn using the *gather_draws* and *add_predicted_draws* functions of the package *tidybayes*^106^. For all models an r-hat of <1.05 was observed indicating that between– and within-chain variance was equal, and thus chains had converged in their parameter estimates. Bulk and tail effective sample sizes (ESS) were high in all models; >2000 for all parameters in the phylogenetically controlled within-species analyses, and >500 for all parameters in the ‘all taxa’ analyses.

## Author contributions

J.W. conducted bioinformatic and statistical analyses, conceptualised research, contributed to design of mathematical modelling, and took the lead in writing the manuscript. P.B. conducted mathematical modelling and wrote the respective methods sections. E.P. contributed to bioinformatic data collection. M.S. aided in supervision and conceptualisation of research. M.R. supervised mathematical modelling. S.v.H. and E.W. supervised and conceptualised research and contributed to design of mathematical modelling. All authors provided critical feedback throughout project and contributed to writing of the manuscript.

## Competing interests

The authors declare no financial competing interests.

## Funding

J.W. is funded by a PhD studentship supervised by M.S., S.v.H., and E.W., and was awarded by the Biotechnology and Biological Sciences Research Council’s (BBSRC’s) South West Biosciences Doctoral Training Partnership (SWBio DTP). E.W., S.v.H., M.R., and M.S. received funding from the BBSRC sLoLa BB/X003051/1.

## Data availability

All genomic data used in this article are publicly available from the NCBI’s *RefSeq* database. Data collected on these genomes and all scripts used for mathematical modelling, bioinformatic data collection, and statistical analyses will be published in publicly accessible repositories upon manuscript acceptance.

## Notes

### Competing Interest Statement

The authors have declared no competing interest.

