## Supplementary Information for "Eco-evolutionary feedbacks drive the co-occurrence of restriction-modification systems and antimicrobial resistance genes"

#### S1.0 Supplementary Results

**Table S1.** Statistics output from the Bayesian models utilised in the 'Phylogenetically controlled within-species analyses'. 'Estimate' denotes the median value drawn from the posterior distribution of the effect of the explanatory variable (Effect) on the response variable (ARG count per genome). 'lwr 95% CI' denotes the lower 95% credibility interval of the estimate, 'upr 95% CI' denotes the upper 95% credibility interval of the estimate, R-hat is a quantification of the Markov chain Monte Carlo chain convergence, Bulk ESS is the effective sample size for rank normalized values using split chains, Tail ESS is the minimum of the effective sample sizes for 5% and 95% quantiles.

| Model | Effect | Estimate | lwr 95% CI | upr 95% CI | R-hat | Bulk ESS | Tail ESS |
|---|---|---|---|---|---|---|---|
| *P. aeruginosa* | Intercept | -2.135 | -2.577 | -1.707 | 1.003 | 5,862 | 2,753 |
| *P. aeruginosa* | RM count | 0.05711 | 0.039 | 0.075 | 1.001 | 7,416 | 3,076 |
| *P. aeruginosa* | Genome length (mb) | 0.5892 | 0.531 | 0.647 | 1.002 | 7,828 | 3,079 |
| *P. aeruginosa* | Phylogeny | 0.2691 | 0.246 | 0.293 | 1.001 | 3,603 | 3,486 |
| *A. baumannii* | Intercept | -1.47 | -2.375 | -0.539 | 1.001 | 2,093 | 2,473 |
| *A. baumannii* | RM count | 0.06892 | 0.030 | 0.109 | 1.001 | 4,173 | 3,518 |
| *A. baumannii* | Genome length (mb) | 0.7375 | 0.626 | 0.846 | 1.000 | 4,682 | 3,515 |
| *A. baumannii* | Phylogeny | 0.5305 | 0.494 | 0.568 | 1.001 | 2,067 | 2,958 |
| *E. faecium* | Intercept | -0.5981 | -1.156 | -0.057 | 1.000 | 4,077 | 3,204 |
| *E. faecium* | RM count | 0.04608 | 0.008 | 0.083 | 1.000 | 7,153 | 3,082 |
| *E. faecium* | Genome length (mb) | 0.6829 | 0.546 | 0.822 | 1.001 | 7,128 | 3,007 |
| *E. faecium* | Phylogeny | 0.4891 | 0.441 | 0.541 | 1.000 | 2,022 | 2,825 |
| *S. pyogenes* | Intercept | -9.622 | -12.436 | -6.936 | 1.000 | 5,164 | 3,189 |
| *S. pyogenes* | RM count | 0.9411 | 0.557 | 1.337 | 1.002 | 3,918 | 3,206 |
| *S. pyogenes* | Genome length (mb) | 5.046 | 3.581 | 6.512 | 1.000 | 5,255 | 3,001 |

| Model | Effect | Estimate | lwr 95% CI | upr 95% CI | R-hat | Bulk ESS | Tail ESS |
|---|---|---|---|---|---|---|---|
| *S. pyogenes* | Phylogeny | 2.102 | 1.858 | 2.381 | 1.001 | 1,997 | 2,885 |
| *N. gonorrhoeae* | Intercept | -1.917 | -5.589 | 1.858 | 1.001 | 7,315 | 3,153 |
| *N. gonorrhoeae* | RM count | 0.2002 | -0.148 | 0.561 | 1.000 | 5,863 | 3,578 |
| *N. gonorrhoeae* | Genome length (mb) | 0.1639 | -0.732 | 1.061 | 1.003 | 13,216 | 2,748 |
| *N. gonorrhoeae* | Phylogeny | 2.191 | 1.680 | 2.814 | 1.000 | 2,586 | 3,391 |
| *All taxa ran on species* | Intercept | 0.9793 | 0.032 | 2.110 | 1.008 | 531 | 621 |
| *All taxa ran on species* | RM count | 0.07413 | 0.057 | 0.091 | 1.001 | 1,549 | 1,621 |
| *All taxa ran on species* | Species identity | 2.028 | 1.362 | 3.104 | 1.006 | 486 | 917 |
| *All taxa no ran on species* | Intercept | 2.16 | 2.144 | 2.175 | 1.001 | 4,256 | 3,292 |
| *All taxa no ran on species* | RM count | -0.4218 | -0.433 | -0.411 | 1.001 | 1,175 | 1,487 |

**Table S2.** Shannon diversity indices for the RM system content of each species, and the P value for the Hutcheson t-test conducted to test if the respective species' diversity index is significantly lower than the Shannon diversity index of all species combined.

| *Species* | *Shannon diversity index* | *P value* |
|---|---|---|
| *Campylobacter jejuni* | 2.178 | > 0.001 |
| *Helicobacter pylori* | 2.897 | > 0.001 |
| *Pseudomonas aeruginosa* | 2.238 | > 0.001 |
| *Acinetobacter baumannii* | 0.943 | > 0.001 |
| *Neisseria gonorrhoeae* | 2.284 | > 0.001 |
| *Klebsiella pneumoniae* | 2.298 | > 0.001 |
| *Shigella flexneri* | 0.4 | > 0.001 |

| *Species* | *Shannon diversity index* | *P value* |
| --- | --- | --- |
| *Shigella sonnei* | 0.638 | > 0.001 |
| *Staphylococcus aureus* | 1.489 | > 0.001 |
| *Streptococcus pyogenes* | 0.374 | > 0.001 |
| *Enterococcus faecium* | 1.258 | > 0.001 |
| *Clostridioides difficile* | 1.276 | > 0.001 |
| *Mycobacterium tuberculosis* | 0.689 | > 0.001 |
| *Salmonella enterica* | 1.936 | > 0.001 |
| *All species combined* | 3.44 | |

16

**Table S3.** The pairwise probabilities that an HGT event between a donor of one species (rows) and a recipient of another species (columns) will be restricted by RM (due to the recipient having an RM system that the donor does not). Values indicate the mean probabilities for all RM systems present in the recipient species. 'Conspecific donor' indicates that the donor is the same species as the recipient.

| *Recipient →*<br>*Donor ↓* | *Acinetobacter baumannii* | *Campylobacter jejuni* | *Clostridioides difficile* | *Enterococcus faecium* | *Helicobacter pylori* | *Klebsiella pneumoniae* | *Mycobacterium tuberculosis* | *Neisseria gonorrhoeae* | *Pseudomonas aeruginosa* | *Salmonella enterica* | *Shigella flexneri* | *Shigella sonnei* | *Staphylococcus aureus* | *Streptococcus pyogenes* |
|---|---|---|---|---|---|---|---|---|---|---|---|---|---|---|
| *Acinetobacter baumannii* | | 0.942 | 0.864 | 0.370 | 0.999 | 0.656 | 1.000 | 1.000 | 0.601 | 0.990 | 0.978 | 0.978 | 0.929 | 0.973 |
| *Campylobacter jejuni* | 0.926 | | 0.985 | 0.932 | 0.995 | 0.963 | 1.000 | 0.999 | 0.957 | 0.998 | 0.998 | 0.998 | 0.992 | 0.996 |
| *Clostridioides difficile* | 0.857 | 0.988 | | 0.870 | 1.000 | 0.929 | 1.000 | 1.000 | 0.917 | 0.998 | 0.995 | 0.995 | 0.985 | 0.994 |
| *Enterococcus faecium* | 0.369 | 0.946 | 0.875 | | 0.999 | 0.685 | 1.000 | 1.000 | 0.635 | 0.991 | 0.980 | 0.980 | 0.935 | 0.975 |
| *Helicobacter pylori* | 0.951 | 0.991 | 0.990 | 0.955 | | 0.975 | 0.999 | 0.991 | 0.971 | 0.993 | 0.998 | 0.998 | 0.995 | 0.994 |
| *Klebsiella pneumoniae* | 0.651 | 0.970 | 0.931 | 0.681 | 0.999 | | 1.000 | 0.999 | 0.798 | 0.994 | 0.988 | 0.989 | 0.964 | 0.985 |
| *Mycobacterium tuberculosis* | 0.988 | 0.999 | 0.998 | 0.989 | 0.999 | 0.994 | | 1.000 | 0.993 | 0.997 | 1.000 | 1.000 | 0.999 | 1.000 |
| *Neisseria gonorrhoeae* | 0.944 | 0.994 | 0.989 | 0.949 | 0.991 | 0.971 | 1.000 | | 0.967 | 0.992 | 0.998 | 0.998 | 0.994 | 0.992 |
| *Pseudomonas aeruginosa* | 0.597 | 0.966 | 0.920 | 0.631 | 0.999 | 0.799 | 1.000 | 1.000 | | 0.994 | 0.986 | 0.987 | 0.958 | 0.984 |
| *Salmonella enterica* | 0.975 | 0.997 | 0.995 | 0.977 | 0.994 | 0.987 | 0.997 | 0.993 | 0.985 | | 0.998 | 0.999 | 0.997 | 0.999 |
| *Shigella flexneri* | 0.971 | 0.998 | 0.994 | 0.974 | 1.000 | 0.985 | 1.000 | 1.000 | 0.983 | 0.999 | | 0.999 | 0.997 | 0.999 |
| *Shigella sonnei* | 0.971 | 0.998 | 0.994 | 0.974 | 1.000 | 0.986 | 1.000 | 1.000 | 0.983 | 1.000 | 0.999 | | 0.997 | 0.999 |
| *Staphylococcus aureus* | 0.921 | 0.993 | 0.984 | 0.928 | 1.000 | 0.961 | 1.000 | 1.000 | 0.954 | 0.999 | 0.997 | 0.997 | | 0.997 |
| *Streptococcus pyogenes* | 0.966 | 0.996 | 0.993 | 0.969 | 0.996 | 0.982 | 1.000 | 0.994 | 0.980 | 0.999 | 0.999 | 0.999 | 0.997 | |
| *Conspecific donor* | 0.310 | 0.986 | 0.967 | 0.423 | 0.971 | 0.825 | 0.987 | 0.939 | 0.766 | 0.990 | 0.992 | 0.992 | 0.987 | 0.991 |

Probability recipient has RM donor does not

0.00 0.25 0.50 0.75 1.00

**Table S4**. Jaccard indices of the ARG repertoire for each pair of species in the dataset. Values fall between 0 and 1, where 0 indicates the respective species share no ARGs, and 1 indicates both species possess an identical repertoire of ARGs.

| | *Acinetobacter baumannii* | *Campylobacter jejuni* | *Clostridioides difficile* | *Enterococcus faecium* | *Helicobacter pylori* | *Klebsiella pneumoniae* | *Mycobacterium tuberculosis* | *Neisseria gonorrhoeae* | *Pseudomonas aeruginosa* | *Salmonella enterica* | *Shigella flexneri* | *Shigella sonnei* | *Staphylococcus aureus* | *Streptococcus pyogenes* |
|---|---|---|---|---|---|---|---|---|---|---|---|---|---|---|
| *Acinetobacter baumannii* | | | | | | | | | | | | | | |
| *Campylobacter jejuni* | 0.007 | | | | | | | | | | | | | |
| *Clostridioides difficile* | 0.011 | 0.146 | | | | | | | | | | | | |
| *Enterococcus faecium* | 0.011 | 0.114 | 0.160 | | | | | | | | | | | |
| *Helicobacter pylori* | 0.013 | 0.050 | 0.070 | 0.021 | | | | | | | | | | |
| *Klebsiella pneumoniae* | 0.129 | 0.019 | 0.028 | 0.044 | 0.016 | | | | | | | | | |
| *Mycobacterium tuberculosis* | 0.010 | 0.015 | 0.049 | 0.031 | 0.133 | 0.010 | | | | | | | | |
| *Neisseria gonorrhoeae* | 0.005 | 0.017 | 0.035 | 0.021 | 0.091 | 0.010 | 0.071 | | | | | | | |
| *Pseudomonas aeruginosa* | 0.117 | 0.011 | 0.016 | 0.017 | 0.013 | 0.189 | 0.008 | 0.008 | | | | | | |
| *Salmonella enterica* | 0.136 | 0.023 | 0.042 | 0.036 | 0.023 | 0.335 | 0.017 | 0.024 | 0.186 | | | | | |
| *Shigella flexneri* | 0.081 | 0.008 | 0.017 | 0.013 | 0.014 | 0.130 | 0.014 | 0.015 | 0.080 | 0.275 | | | | |
| *Shigella sonnei* | 0.104 | 0.007 | 0.022 | 0.023 | 0.022 | 0.184 | 0.021 | 0.045 | 0.113 | 0.360 | 0.411 | | | |
| *Staphylococcus aureus* | 0.017 | 0.090 | 0.101 | 0.282 | 0.050 | 0.032 | 0.048 | 0.038 | 0.034 | 0.042 | 0.029 | 0.038 | | |
| *Streptococcus pyogenes* | 0.010 | 0.125 | 0.197 | 0.172 | 0.067 | 0.023 | 0.061 | 0.154 | 0.015 | 0.032 | 0.035 | 0.046 | 0.214 | |

Jaccard indices of ARGs

0.00 0.25 0.50 0.75 1.00

**Table S5**. Pairs of RM systems and ARGs that co-occur significantly more than chance (using Bonferroni corrected alpha values), where the effect size of the co-occurrence is large (observed co-occurrence >= 10* expected co-occurrence) and excluding rare pairs that co-occur in fewer than 0.5% of the respective species' genomes.

| Species | RM hmm name | ARG name | RM system n | ARG n | obs/exp | P | ARG description | resistance | mean proximity (bp) | proportion on same contig |
|---|---|---|---|---|---|---|---|---|---|---|
| *S. pyogenes* | REase_II_00151 | mef(A) | 19 | 19 | Inf | 7.41e-29 | macrolide efflux MFS transporter Mef(A) | macrolide | 9,257 | 1.00 |
| *S. pyogenes* | REase_II_00151 | msr(D) | 19 | 19 | Inf | 7.41e-29 | ABC-F type ribosomal protection protein Msr(D) | macrolide | 7,756 | 1.00 |
| *P. aeruginosa* | REase_I_00002 | dfrB5 | 84 | 84 | 46.00 | 2.71e-51 | trimethoprim-resistant dihydrofolate reductase DfrB5 | trimethoprim | 1,005,859 | 0.26 |
| *P. aeruginosa* | REase_I_00002 | blaOXA-4 | 84 | 51 | 45.00 | 2.64e-59 | carbapenem-hydrolyzing class D beta-lactamase OXA-4 | carbapenem | 1,648,450 | 0.16 |
| *P. aeruginosa* | REase_I_00002 | aac(3)-Id | 84 | 75 | 44.00 | 2.07e-50 | aminoglycoside N-acetyltransferase AAC(3)-Id | gentamicin | 1,090,488 | 0.25 |
| *A. baumanii* | REase_I_00005 | blaADC-182 | 80 | 52 | 40.00 | 1.97e-48 | class C beta-lactamase ADC-182 | cephalosporin | 388,574 | 0.22 |
| *P. aeruginosa* | REase_I_00003 | blaPDC-36 | 59 | 42 | 40.00 | 3.04e-60 | class C beta-lactamase PDC-36 | cephalosporin | 1,918,323 | 0.12 |
| *P. aeruginosa* | REase_I_00002 | cmlA6 | 84 | 51 | 40.00 | 6.82e-51 | chloramphenicol efflux MFS transporter CmlA6 | chloramphenicol | 2,100,532 | 0.02 |
| *A. baumanii* | REase_I_00005 | blaTEM-150 | 80 | 41 | 34.00 | 1.72e-42 | class A beta-lactamase TEM-150 | beta-lactam | 2,060,794 | 0.09 |
| *P. aeruginosa* | REase_MTase_IIG_00002 | blaVEB-9 | 155 | 40 | 29.00 | 6.60e-29 | class A extended-spectrum beta-lactamase VEB-9 | cephalosporin | 158,144 | 0.48 |
| *A. baumanii* | REase_I_00005 | blaOXA-51 | 80 | 56 | 27.00 | 6.59e-28 | OXA-51 family carbapenem-hydrolyzing class D beta-lactamase OXA-51 | carbapenem | 853,974 | 0.04 |
| *P. aeruginosa* | REase_MTase_IIG_00002 | dfrB2 | 155 | 30 | 26.00 | 5.14e-28 | trimethoprim-resistant dihydrofolate reductase DfrB2 | trimethoprim | 144,640 | 0.54 |
| *P. aeruginosa* | REase_I_00002 | aph(3')-XV | 84 | 64 | 25.00 | 4.46e-25 | aminoglycoside O-phosphotransferase APH(3')-XV | amikacin kanamycin | 5,298,421 | 0.16 |
| *P. aeruginosa* | REase_II_00142 | aac(6')-29a | 225 | 51 | 24.50 | 7.04e-46 | aminoglycoside N-acetyltransferase AAC(6')-29a | aminoglycoside | 1,293,581 | 0.04 |
| *P. aeruginosa* | REase_I_00002 | aadA2 | 84 | 137 | 22.50 | 1.24e-40 | ANT(3'')-Ia family aminoglycoside nucleotidyltransferase AadA2 | streptomycin | 975,494 | 0.27 |
| *P. aeruginosa* | REase_I_00009 | blaOXA-56 | 281 | 42 | 20.00 | 5.94e-34 | OXA-10 family oxacillin-hydrolyzing class D beta-lactamase OXA-56 | beta-lactam | 3,737,765 | 0.15 |
| *P. aeruginosa* | REase_I_00002 | tet(G) | 84 | 192 | 19.33 | 6.64e-50 | tetracycline efflux MFS transporter Tet(G) | tetracycline | 3,155,011 | 0.28 |
| *P. aeruginosa* | REase_II_00142 | blaOXA-9 | 225 | 38 | 19.00 | 1.42e-36 | oxacillin-hydrolyzing class D beta-lactamase OXA-9 | beta-lactam | 3,154,473 | 0.28 |
| *P. aeruginosa* | REase_I_00002 | floR2 | 84 | 177 | 19.00 | 1.44e-50 | chloramphenicol/florfenicol efflux MFS transporter FloR2 | chloramphenicol florfenicol | 3,229,409 | 0.08 |
| *P. aeruginosa* | REase_II_00142 | cmlB | 225 | 37 | 18.50 | 1.18e-35 | chloramphenicol efflux MFS transporter CmlB1 | chloramphenicol | 3,594,717 | 0.05 |
| *A. baumanii* | REase_I_00005 | dfrA1 | 80 | 100 | 18.00 | 2.52e-32 | trimethoprim-resistant dihydrofolate reductase DfrA14 | trimethoprim | 2,062,830 | 0.08 |

| Species | RM hmm name | ARG name | RM system n | ARG n | obs/exp | P | ARG description | resistance | mean proximity (bp) | proportion on same contig |
|---|---|---|---|---|---|---|---|---|---|---|
| *A. baumanii* | REase_I_00005 | sat2_gen | 80 | 92 | 18.00 | 1.46e-33 | streptothricin N-acetyltransferase Sat2 | streptothricin | 2,062,606 | 0.08 |
| *P. aeruginosa* | REase_II_00142 | aac(6')-29b | 225 | 56 | 17.67 | 3.68e-49 | aminoglycoside N-acetyltransferase AAC(6')-29b | aminoglycoside | 2,566,367 | 0.02 |
| *P. aeruginosa* | REase_MTase_IIG_00002 | blaOXA-846 | 155 | 108 | 16.50 | 4.67e-59 | OXA-50 family oxacillin-hydrolyzing class D beta-lactamase OXA-846 | beta-lactam | 1,040,772 | 0.76 |
| *P. aeruginosa* | REase_I_00009 | rmtD1 | 281 | 31 | 15.00 | 4.62e-26 | 16S rRNA (guanine(1405)-N(7))-methyltransferase RmtD1 | aminoglycoside | 3,831,774 | 0.13 |
| *P. aeruginosa* | REase_I_00009 | blaSPM-1 | 281 | 28 | 14.00 | 8.33e-25 | subclass B1 metallo-beta-lactamase SPM-1 | carbapenem | 3,630,133 | 0.25 |
| *P. aeruginosa* | REase_I_00009 | blaVEB-9 | 281 | 40 | 13.00 | 1.26e-18 | class A extended-spectrum beta-lactamase VEB-9 | cephalosporin | 2,312,364 | 0.58 |
| *P. aeruginosa* | REase_II_00142 | blaCARB-2 | 225 | 76 | 12.75 | 3.42e-40 | PSE family carbenicillin-hydrolyzing class A beta-lactamase CARB-2 | beta-lactam | 786,362 | 0.06 |
| *P. aeruginosa* | REase_I_00009 | aac(3)-Ic | 281 | 36 | 12.50 | 1.26e-18 | aminoglycoside N-acetyltransferase AAC(3)-Ic | gentamicin | 2,040,155 | 0.16 |
| *A. baumanii* | REase_I_00009 | blaADC-191 | 187 | 37 | 12.50 | 1.09e-20 | class C beta-lactamase ADC-191 | cephalosporin | 2,676,636 | 0.04 |
| *A. baumanii* | REase_I_00008 | blaADC-222 | 237 | 30 | 12.00 | 2.91e-19 | class C beta-lactamase ADC-222 | cephalosporin | 757,849 | 0.08 |
| *A. baumanii* | REase_I_00008 | blaOXA-95 | 237 | 29 | 12.00 | 1.36e-19 | OXA-51 family carbapenem-hydrolyzing class D beta-lactamase OXA-95 | carbapenem | 2,146,325 | 0.08 |
| *A. baumanii* | REase_I_00008 | blaADC-26 | 237 | 108 | 11.83 | 2.62e-48 | class C extended-spectrum beta-lactamase ADC-26 | cephalosporin | 2,096,475 | 0.07 |
| *P. aeruginosa* | REase_MTase_IIG_00003 | blaPDC-34 | 397 | 183 | 11.80 | 1.85e-121 | class C beta-lactamase PDC-34 | cephalosporin | 800,446 | 0.08 |
| *P. aeruginosa* | REase_I_00002 | aac(6')-II | 84 | 213 | 11.75 | 9.15e-35 | aminoglycoside N-acetyltransferase AAC(6')-II | amikacin;kanamycin tobramycin | 1,006,579 | 0.26 |
| *A. baumanii* | REase_I_00005 | blaOXA-65 | 80 | 129 | 11.33 | 2.65e-26 | OXA-51 family carbapenem-hydrolyzing class D beta-lactamase OXA-65 | carbapenem | 786,501 | 0.03 |
| *P. aeruginosa* | REase_I_00002 | blaVIM-2 | 84 | 231 | 11.25 | 4.04e-31 | subclass B1 metallo-beta-lactamase VIM-2 | carbapenem | 594,149 | 0.16 |
| *P. aeruginosa* | REase_MTase_IIG_00002 | tet(A) | 155 | 84 | 11.00 | 2.04e-24 | tetracycline efflux MFS transporter Tet(A) | tetracycline | 155,749 | 0.36 |
| *P. aeruginosa* | REase_III_00001 | blaGES-14 | 347 | 36 | 10.33 | 9.01e-23 | carbapenem-hydrolyzing class A beta-lactamase GES-14 | carbapenem | 20,557 | 0.19 |
| *P. aeruginosa* | REase_I_00009 | aadA7 | 281 | 60 | 10.25 | 2.86e-29 | ANT(3'')-Ia family aminoglycoside nucleotidyltransferase AadA7 | streptomycin | 3,737,764 | 0.15 |
| *A. baumanii* | REase_I_00005 | cmlB1 | 80 | 176 | 10.25 | 2.31e-29 | chloramphenicol efflux MFS transporter CmlB1 | chloramphenicol | 899,322 | 0.02 |

### S2.0 Supplementary Materials & Methods

#### S2.1 Mathematical modelling

Our mathematical model simulates population dynamics with MGEs being acquired on the basis of the relative rate of HGT and the presence of RM systems. We considered three RM system investment strategies – zero, low, and high. S(t) denotes the rate of change of a sub-population with 'n' MGEs and having RM system 'i', given by the following ODE -

$$\frac{dS_{n,i}(t)}{dt} = \left(\beta\left(1 - c_{mge}\right)^{i}(1 - c_n)\right) S_{n,i}(t)\left(1 - \frac{\sum S(t)}{K}\right) - \varphi S_{n,i}(t)$$

$$- \alpha_h \epsilon \left(\frac{\sum_{n'=n}\sum_{i=1}^{m} S_{n',i}(t)}{T}\right) S_{n,i}(t) - \alpha_n \epsilon \left(\frac{\sum_{n'\neq n}\sum_{i=1}^{m} S_{n',i}(t)}{T}\right) S_{n,i}(t)$$

$$+ i\lambda S_{n,i+1}(t)$$

Individuals are born at rate '$\beta$', scaled by the logistic growth constraint, and die at rate '$\varphi$'. Populations that contain MGEs suffer a reduced birth rate given by $(1 – c_{mge})^m$ where '$c_{mge}$' is the cost of an MGE and 'i' is the number of MGEs present. Populations with investment in an RM system suffer a reduced birth rate given by $(1 – c_n)$ where '$c_n$' is the cost of the RM investment strategy (none, low and high). The acquisition of MGEs depended on the relative rate of HGT given by '$\alpha_h$', the contact factor '$\epsilon$' and the prevalence of MGE carrying cells in the system. The presence of an RM system reduced the chance of acquiring an MGE. However, RM systems are blind towards MGEs that originate from populations with the same RM system, as these MGEs would be methylated. Thus, the acquisition of MGEs by populations with an RM was divided into two components. MGEs from populations with no RM and a different RM system were acquired at a reduced rate. This rate was given by the amount of HGT '$\alpha_n=\alpha_h/R$' which was less than '$\alpha_h$' (by a factor of $2^1$ for low investment RM and a factor of $2^7$ for high investment RM). Acquisition of MGEs from populations with the same RM system did not suffer a reduction in rate, but there was a variation in the prevalence of such populations. Populations with MGEs could also lose MGEs at a rate given by the intrinsic MGE clearance rate '$\lambda$' and the number of MGEs present, given by '*i*'. We modelled the acquisition of up to five MGEs.

For our initial condition, we considered one individual of each phenotype and simulated population growth under varying relative rates of HGT, cost of low and high RM system investment strategies and the cost of MGEs. We simulated the system for four-thousand-time steps, which was sufficient to reach steady state or oscillatory behaviour. See Table S6 for parameters.

**Table S6.** The parameters and the symbols used to denote them in the equations, and the values that were used for them.

| Parameter | Symbol | Value(s) |
|---|---|---|
| Birth rate | b | 2 |
| Carrying capacity | K | 1000 |
| Contact factor | β | 0.9 |
| Cost of one MGE | $c_{mge}$ | 5, 10, 15 and 20% of birth rate |
| Cost of low investment RM | $c_r$ | 0.016, 0.047, 0.093, 0.141 and 0.188% of birth rate |
| Cost of high investment RM | $c_R$ | 1, 3, 6, 9 and 12% of birth rate |
| Death rate | d | 0.5 |
| MGE clearance rate | ʎ | 0.015 |
| HGT exposure | $\alpha_n$ | 0 to 1 |

#### S2.2 Trait depth filtered modelling

To calculate the trait depth of both RM systems and ARGs, the function *consentrait_depth* from the R package *castor* was used(1). Phylogenetic trees used in trait depth estimation were the same used in the phylogenetic controlled within-species modelling (see 'Within-species phylogeny construction'). Settings were informed from previous literature(2). Specifically, the minimum fraction of tips in a clade exhibiting a trait, for the clade to be considered positive in the trait, was 0.9, and all positive clades were weighted equally. Single tips exhibiting a trait were included, and the phylogenetic depth was taken to be half the length of their incoming edge. The mean trait depth of each gene was calculated at the level of the species. The trait depth filtered dataset (see Figure S5), included only the RM systems in the upper tertile for trait depth, and ARGs in the lower tertile for trait depth. These data were then used in phylogenetically controlled Bayesian Poisson generalised linear models, in the same was as described in ‘Materials & Methods: Bayesian mixed-effect modelling’.

##### S2.3 Pairwise ARG Jaccard indices calculation and correlation with pairwise probability donors have RM systems recipients do not

To quantify the pairwise similarities for ARG repertoires between different species, we calculated Jaccard indices. For each pair of ARG sets, the Jaccard index was computed as the number of shared ARGs (i.e., the intersection of the two sets) divided by the total number of unique genes present in either set (i.e., the union of the two sets). This metric ranges from 0 (no shared ARGs) to 1 (identical ARG sets), providing a measure of overlap between species-specific gene repertoires.

We wanted to test if there was a correlation between the pairwise Jaccard indices of ARGs (i.e., the similarity in ARG repertoire between any two species), and the probability that during an HGT event between these two species, a recipient will have an RM system the donor does not (i.e., the probability that RM will pose a barrier to HGT). We hypothesised that pairs of species that had a lower probability that RM would act as a barrier to HGT, would have more similar ARG repertoires (i.e., we predicted a negative correlation between these two metrics). To test this hypothesis, we log 10 transformed both metrics so that they would be normally distributed. We added 10% of the lowest non-zero value to all probabilities a donor will have an RM the recipient does not, in order to prevent infinite values from log transforming zeros. We then used the *ggpubr::stat_cor* function to calculate the Pearson correlation coefficient. To ensure our analysis was robust we also calculated the Pearson correlation coefficient with the zeros removed (just 2.7% of all values), and found our conclusions did not differ.


#### Figure legends

**Figure S1.** The average count of MGEs carried by populations with different levels of RM investment. Simulations when population-level RM diversity is either A) low or B) high. Panels are divided into columns to denote simulations with different fixed MGE costs (as a percentage reduction in birth rate per MGE) and divided into rows to denote simulations with different fixed RM system investment costs (as a percentage reduction in birth rate, values denote the costs of the high RM investment strategy, low investment RM system costs are scaled linearly). Each panel includes data from all sub-populations and simulations across the range of relative HGT rates. The central horizontal line indicates the mean, the boxes indicate ± 1 standard deviation (SD) from the mean, the whiskers denote ±2 SD from the mean.

**Figure S2.1.** The mean count of defence systems per genome for each of the 14 species studied.

**Figure S2.2.** The mean count of antimicrobial resistance genes (ARGs) per genome for each of the 14 species studied.

**Figure S3.1.** *Pseudomonas aeruginosa* phylogenetic tree with presence of ARGs and RM systems. ARGs are shown in the left heatmap, whereas RM systems are shown in the right heatmap. For each heatmap, genes are arranged from most to least abundant, moving from left to right. Note, in order to aid visualisation this tree shows a sub-sample of 200 genomes, not all genomes in the dataset. Where more than 10 different RM systems are observed in the species, the 10 most abundant are shown, and where more than 20 different ARGs are observed in the species, the 20 most abundant are shown.

**Figure S3.2.** *Acinetobacter baumanii* phylogenetic tree with presence of ARGs and RM systems. ARGs are shown in the left heatmap, whereas RM systems are shown in the right heatmap. For each heatmap, genes are arranged from most to least abundant, moving from left to right. Note, in order to aid visualisation this tree shows a sub-sample of 200 genomes, not all genomes in the dataset. Where more than 10 different RM systems are observed in the species, the 10 most abundant are shown, and where more than 20 different ARGs are observed in the species, the 20 most abundant are shown.

**Figure S3.3.** *Enterococcus faecium* phylogenetic tree with presence of ARGs and RM systems. ARGs are shown in the left heatmap, whereas RM systems are shown in the right heatmap. For each heatmap, genes are arranged from most to least abundant, moving from left to right. Note, in order to aid visualisation this tree shows a sub-sample of 200 genomes, not all genomes in the dataset. Where more than 10 different RM systems are observed in the species, the 10 most abundant are shown, and where more than 20 different ARGs are observed in the species, the 20 most abundant are shown.

**Figure S3.4.** *Streptococcus pyogenes* phylogenetic tree with presence of ARGs and RM systems. ARGs are shown in the left heatmap, whereas RM systems are shown in the right heatmap. For each heatmap, genes are arranged from most to least abundant, moving from left to right. Note, in order to aid visualisation this tree shows a sub-sample of 200 genomes, not all genomes in the dataset. Where more than 10 different RM systems are observed in the species, the 10 most abundant are shown, and where more than 20 different ARGs are observed in the species, the 20 most abundant are shown.

**Figure S3.5.** *Neisseria gonorrhoea* phylogenetic tree with presence of ARGs and RM systems. ARGs are shown in the left heatmap, whereas RM systems are shown in the right heatmap. For each heatmap, genes are arranged from most to least abundant, moving from left to right. Note, in order to aid visualisation this tree shows a sub-sample of 200 genomes, not all genomes in the dataset.

Where more than 10 different RM systems are observed in the species, the 10 most abundant are shown, and where more than 20 different ARGs are observed in the species, the 20 most abundant are shown.

**Figure S4**. An example of the region of the *Streptococcus pyogenes* prophage Φ1207.3 where two macrolide resistance genes (mef(A) and msr(D)) co-localise with a Type II RM system (MTase_II is the methyltransferase M.SpyI, and REase_II_SU_1 and REase_II_SU_2 are the two subunits of the restriction enzyme).

**Figure S5.** Comparison of models without trait depth filtering (see Figure 3), to those with trait depth filtering "trait depth filtered". Trait depth filtering refers to a model produced from a dataset where only the RM systems in the upper tertile for trait depth, and ARGs in the lower tertile for trait depth are retained (see S1.2 Trait depth filtered modelling). Distributions are draws from posterior distributions of the effect of RM system count on ARG count per genome, after controlling for phylogeny and genome length. The black horizontal line indicates the 95% of draws that fall closest to the mean. Effect size is the effect each additional RM system has on the count of ARGs in a genome. Note X axis scale is not uniform across species.

**Figure S6.** Kernel density of trait depths for RM systems and ARGs per species.

**Figure S7**. The relationship between the pairwise probabilities a recipient has an RM system the donor does not, and the logged pairwise Jaccard indices (a measure of similarity) for ARG repertoire. *R* denotes Pearson's correlation coefficient and *p* denotes the *p* value for the statistical significance of the correlation.
