## Supplementary figures and images for "Eco-evolutionary feedbacks drive the co-occurrence of restriction-modification systems and antimicrobial resistance genes"

### Figure S1

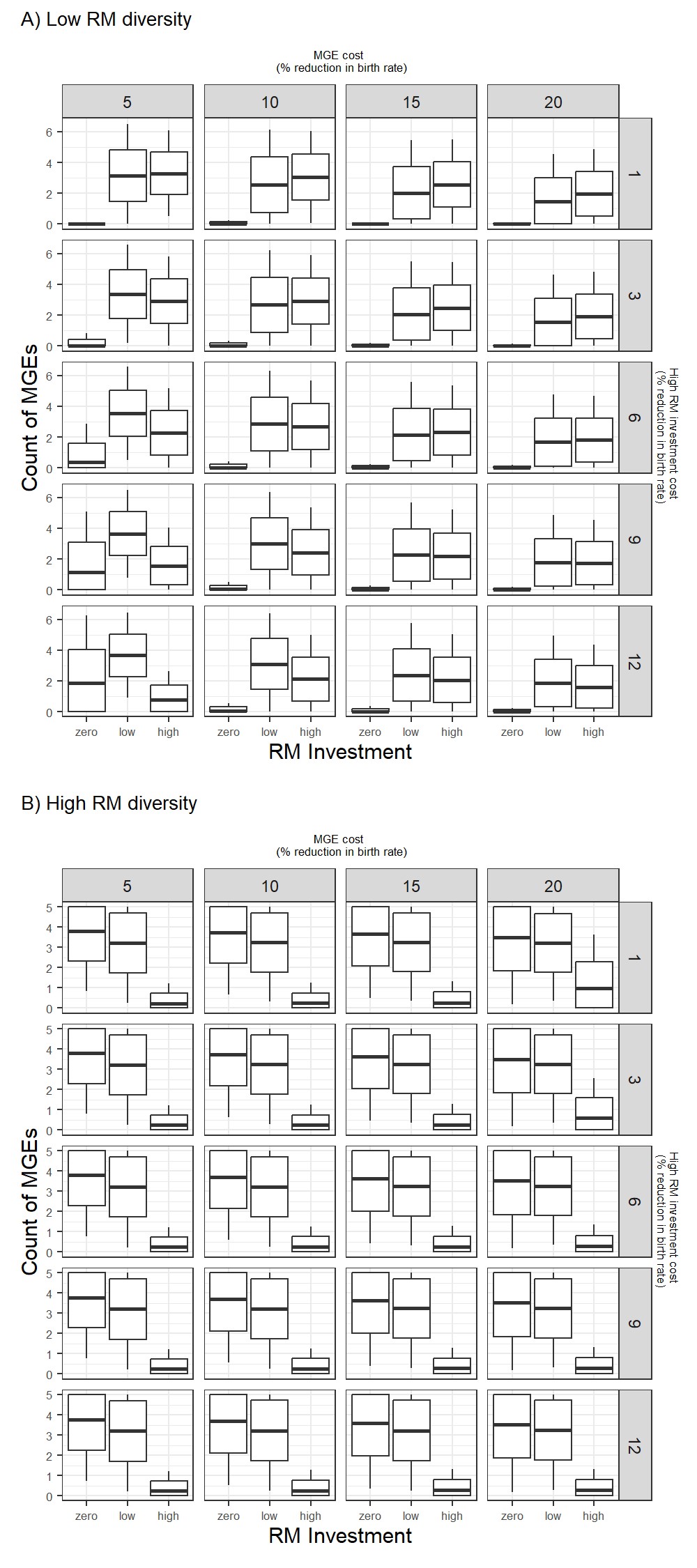

### Figure S2.1

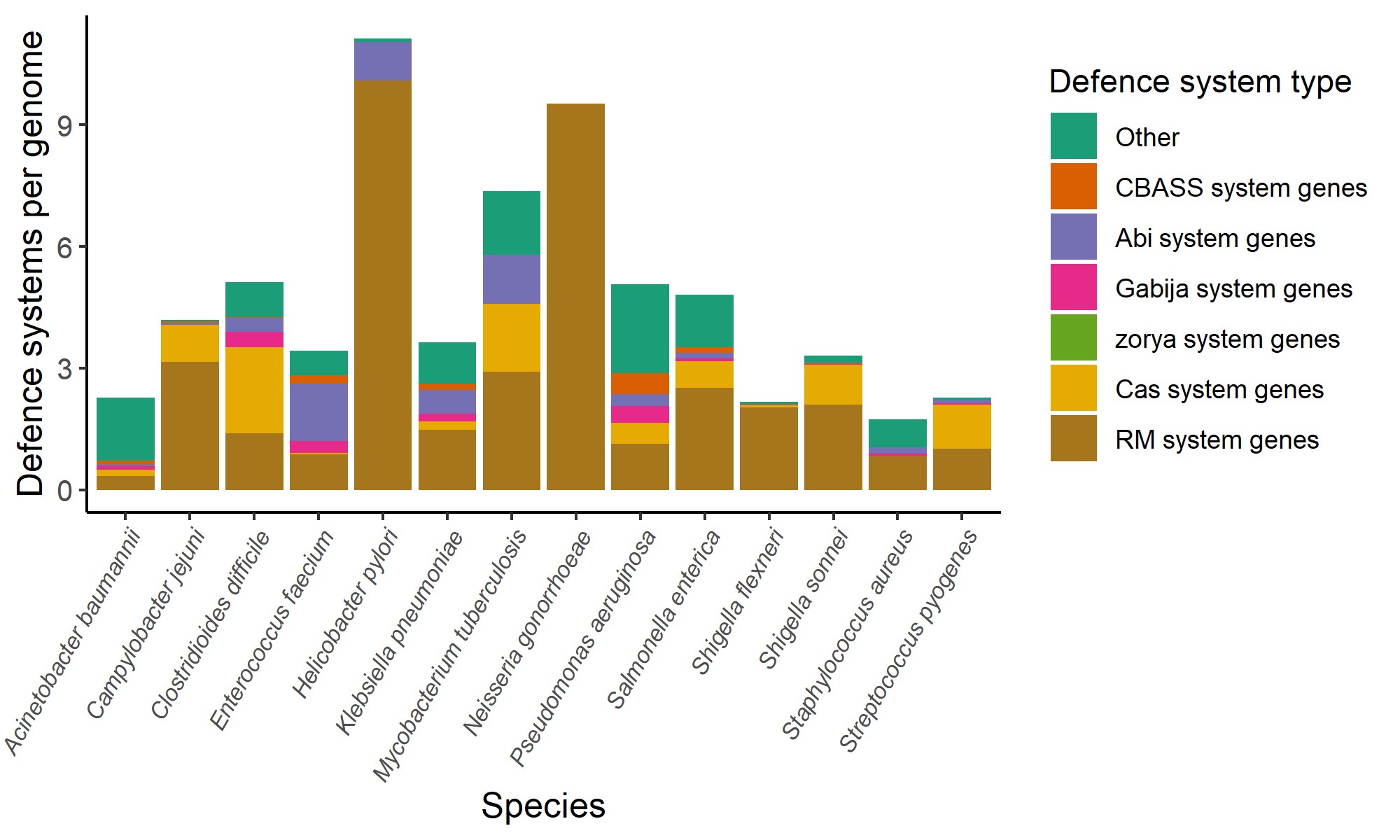

### Figure S2.2

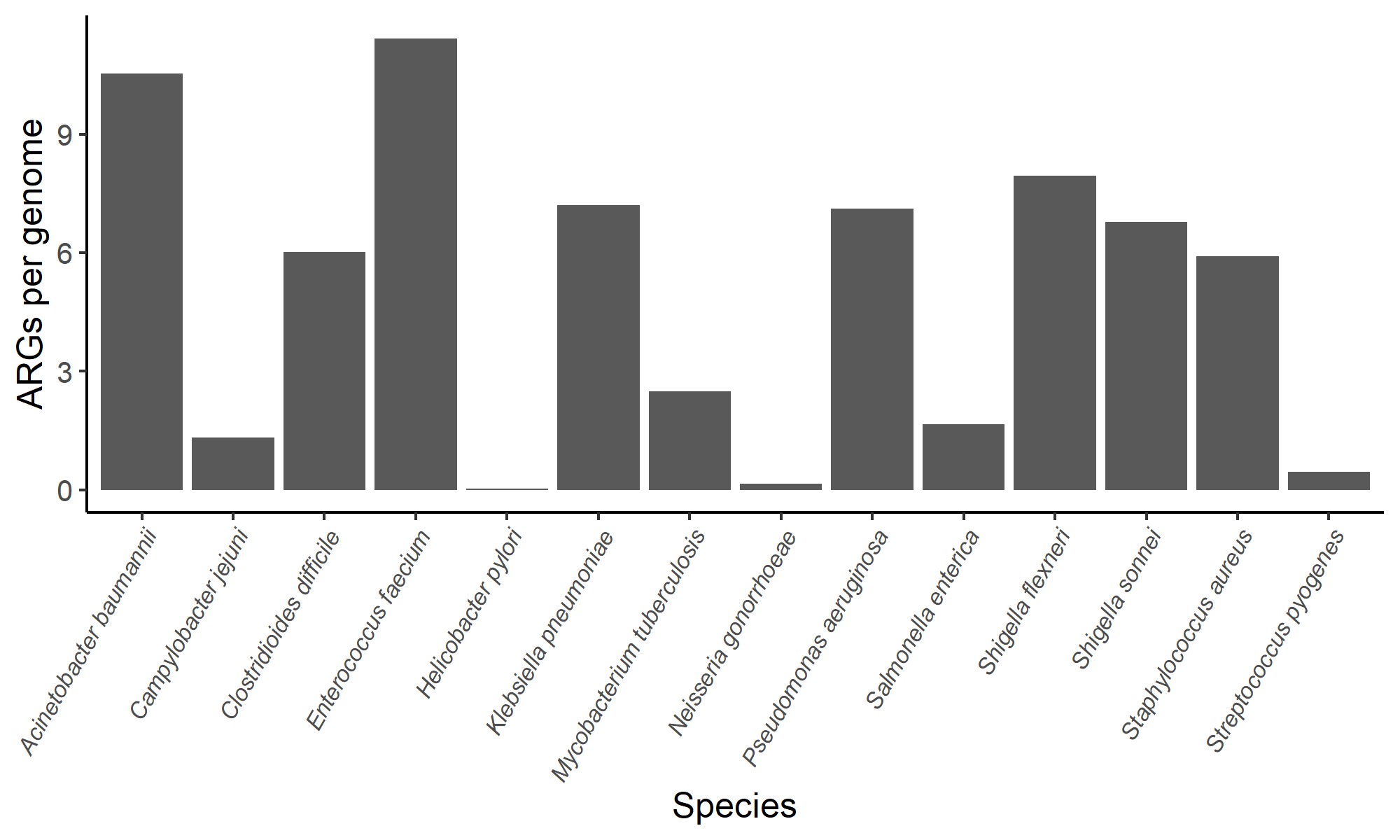

### Figure S3.1

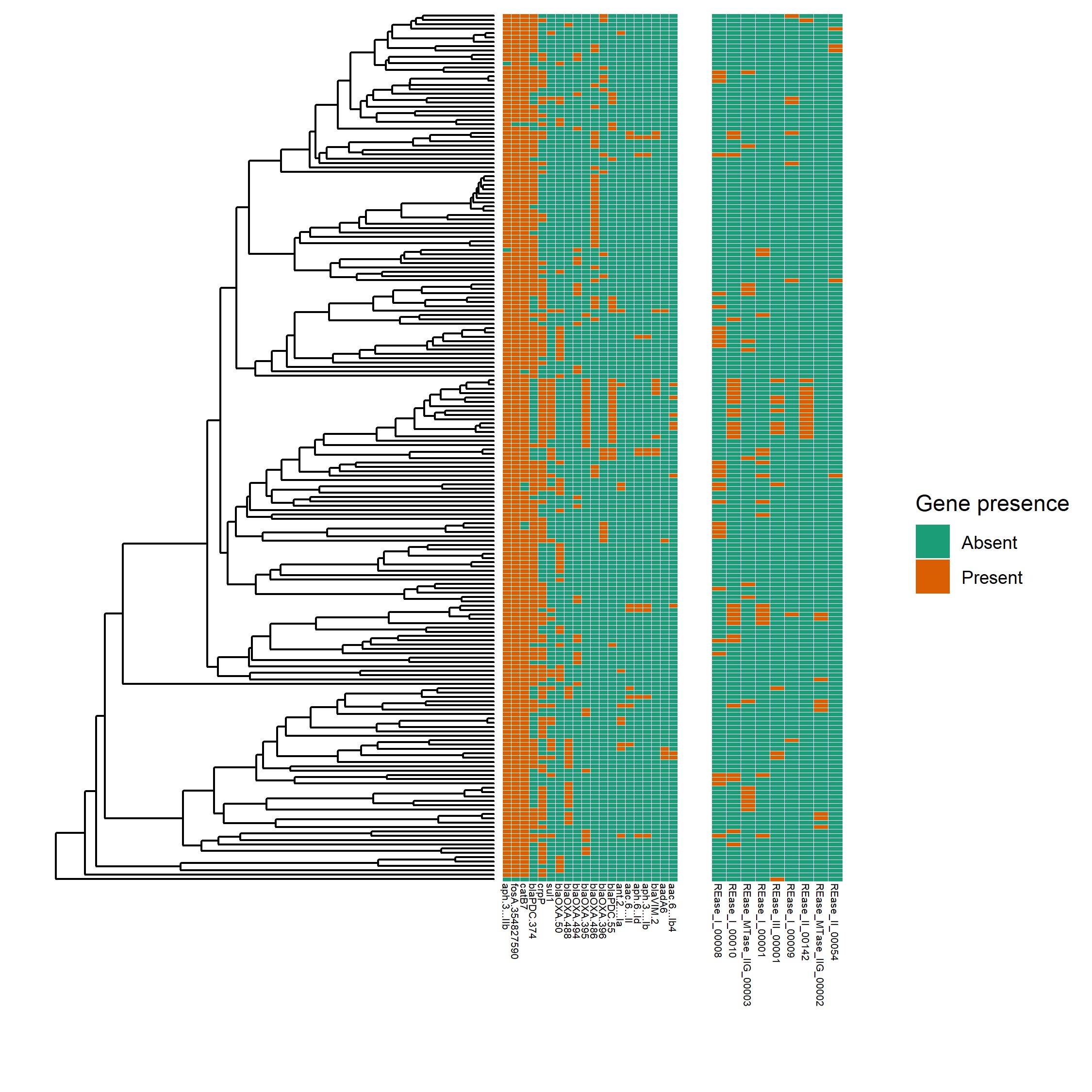

### Figure S3.2

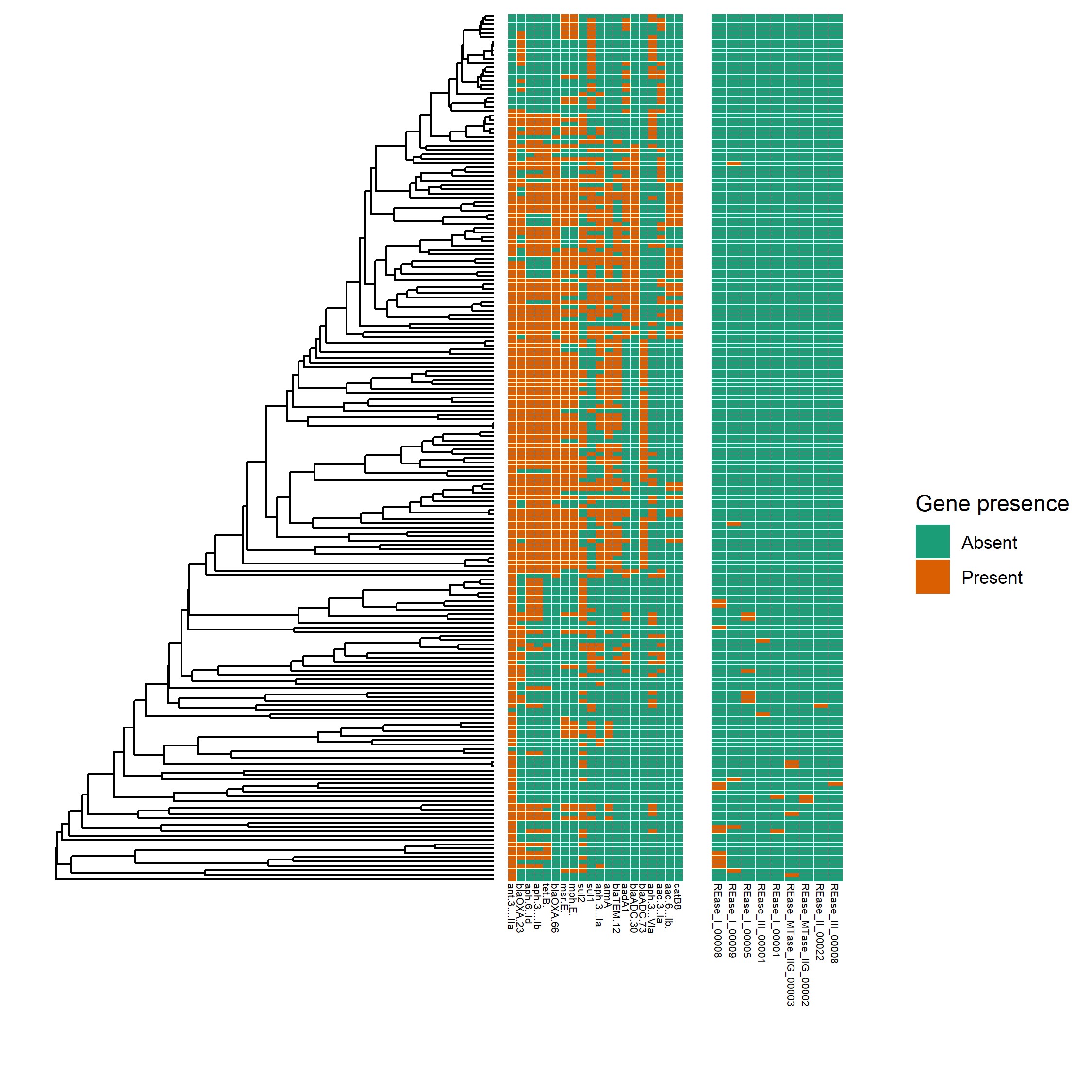

### Figure S3.3

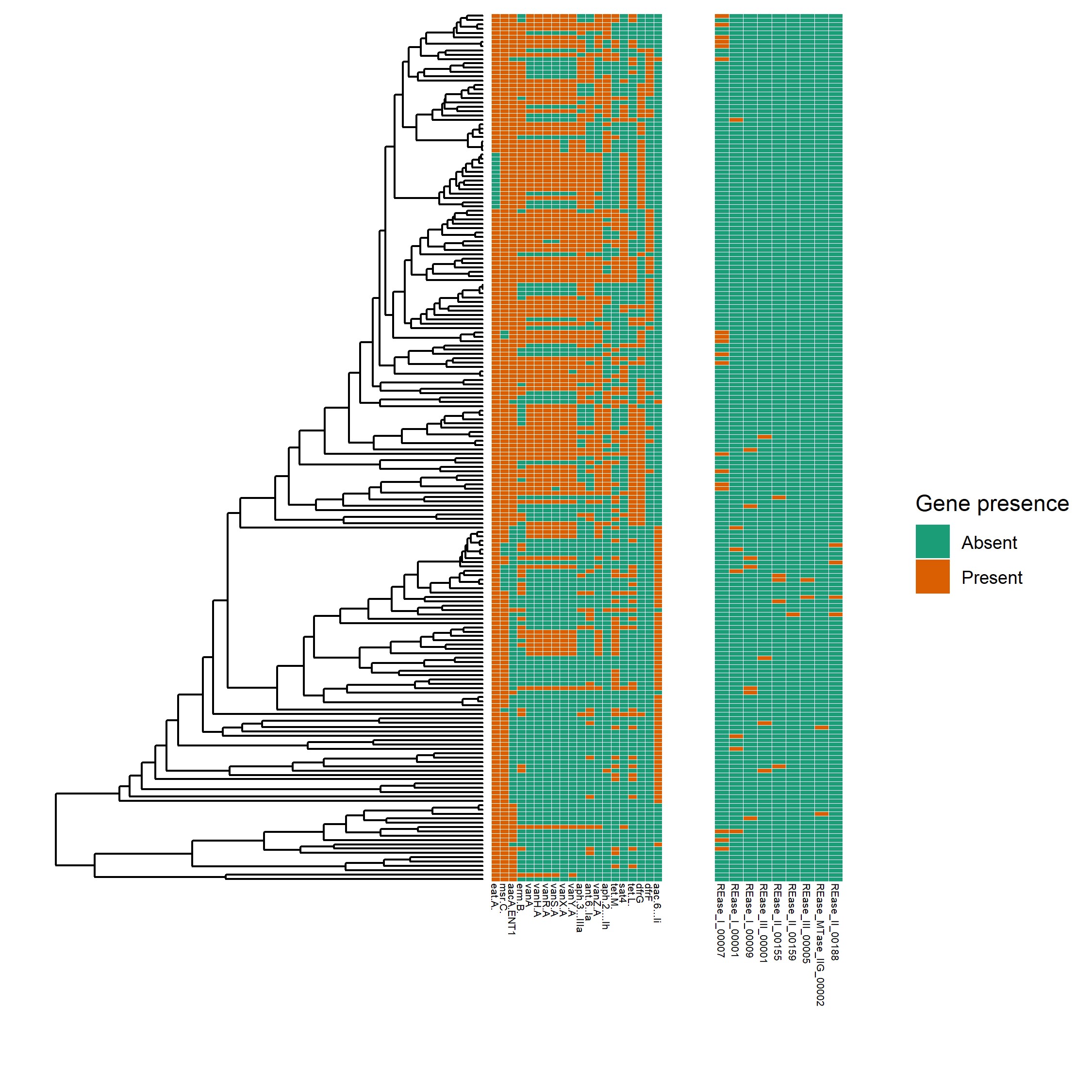

### Figure S3.4

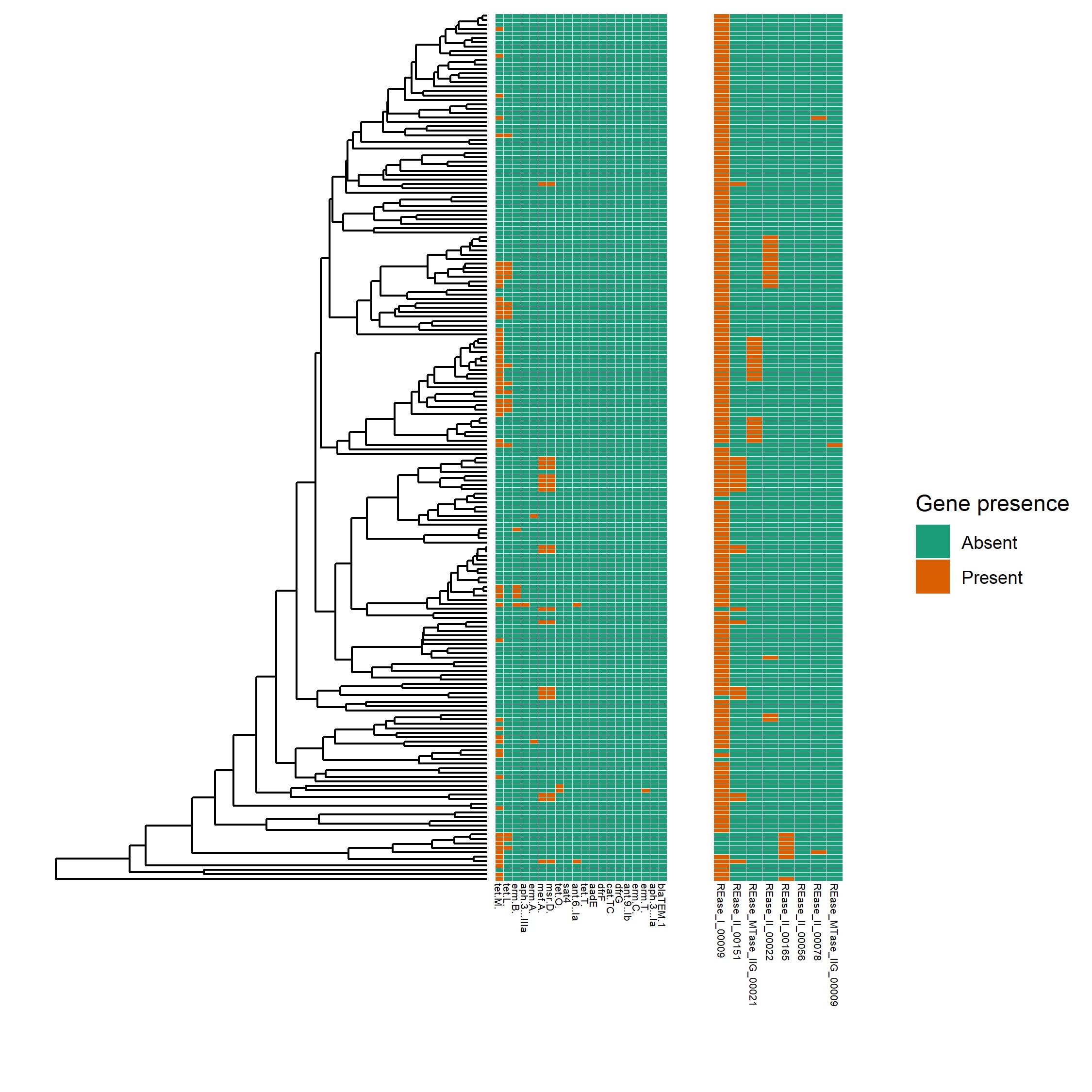

### Figure S3.5

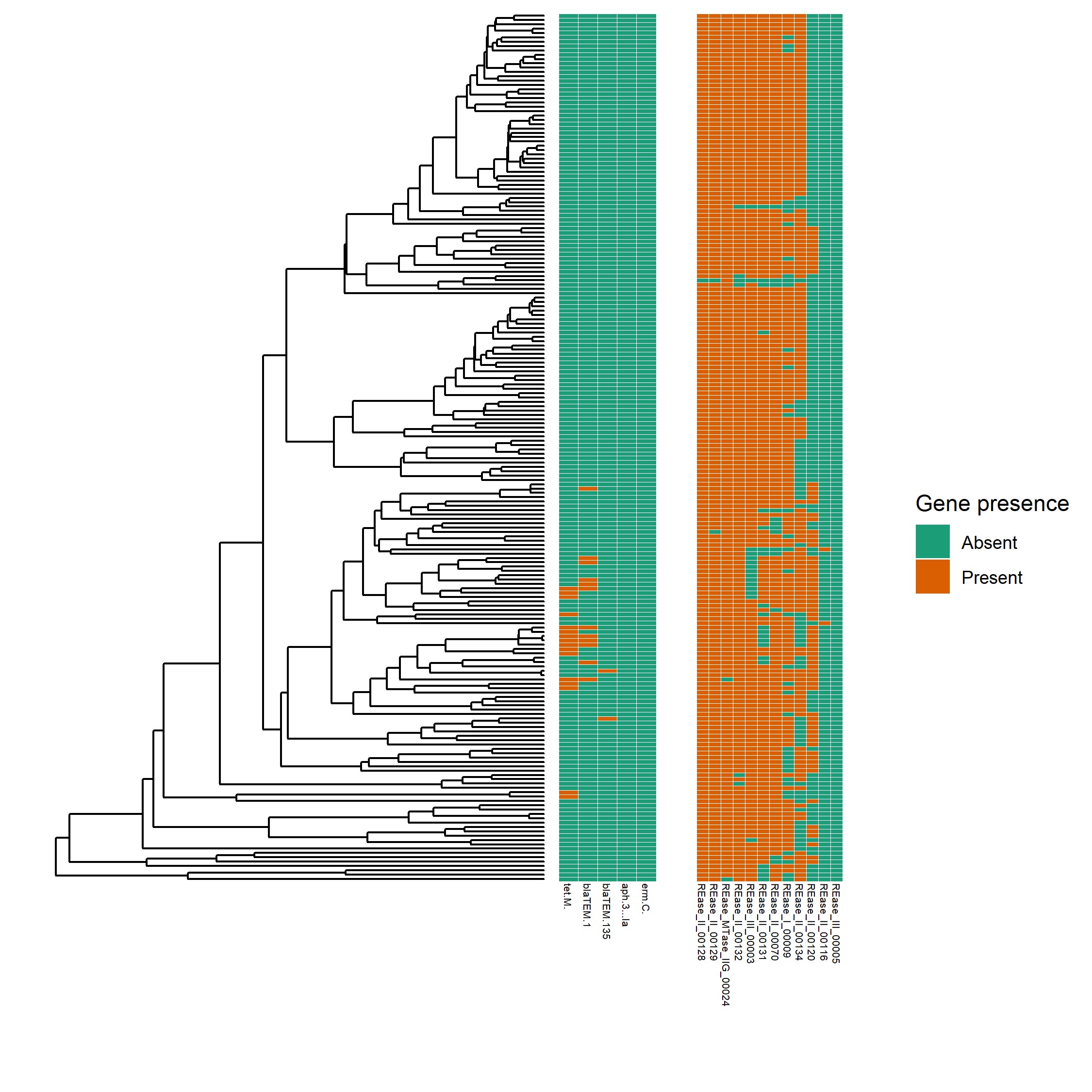

### Figure S4

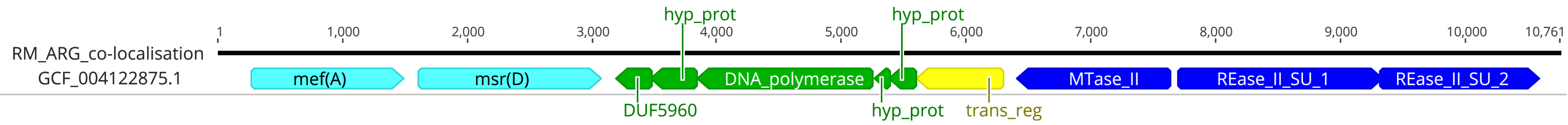

### Figure S5

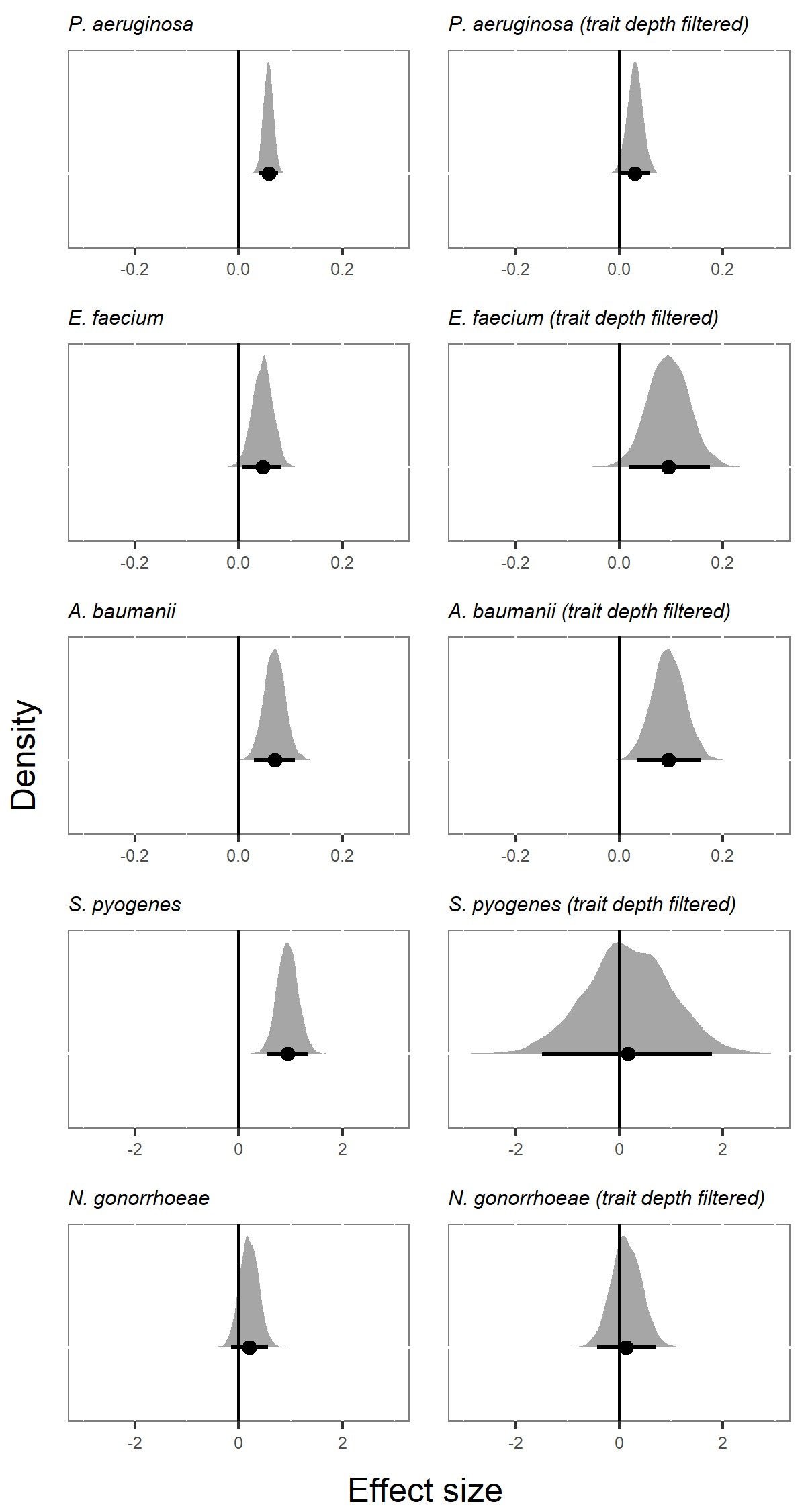

### Figure S6

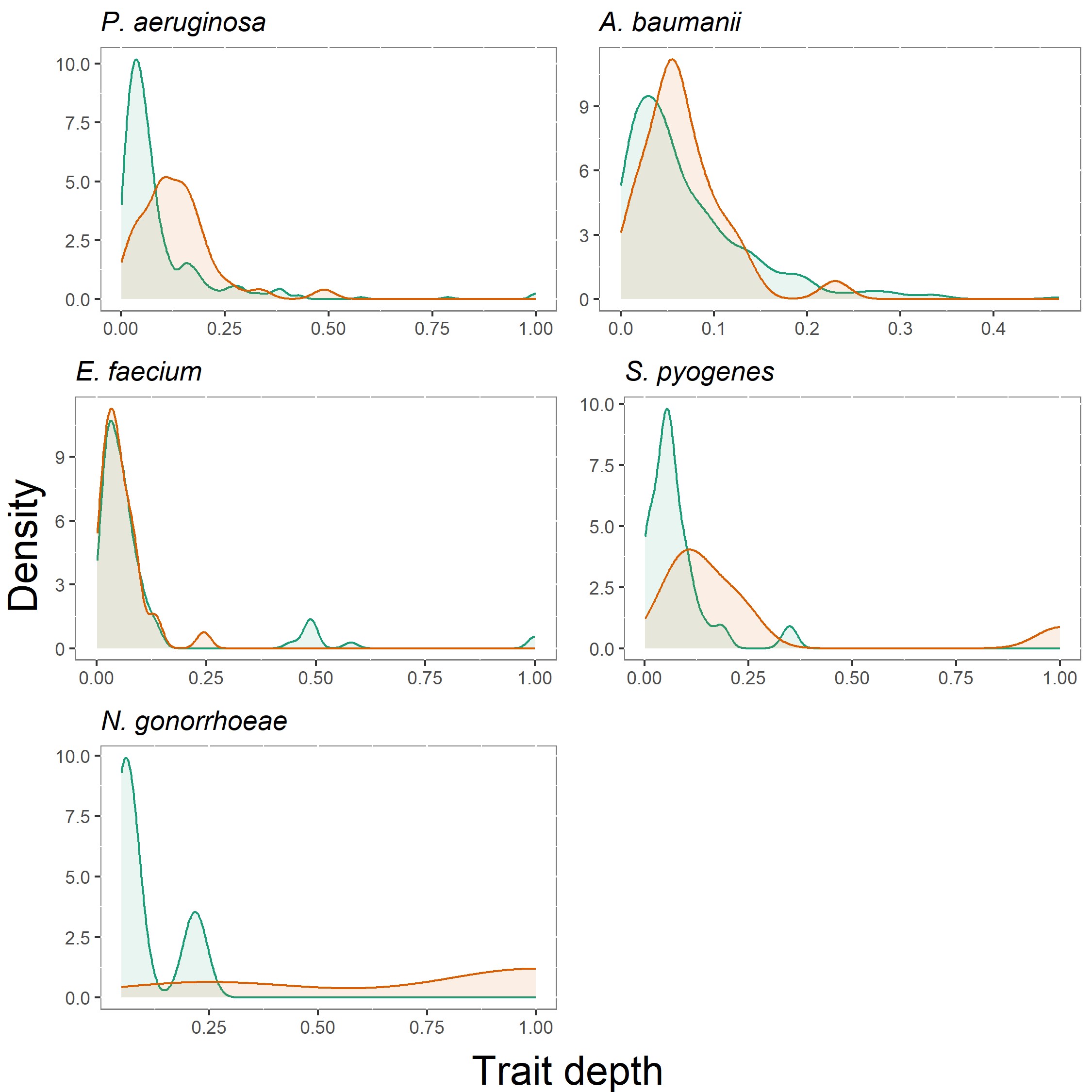

### Figure S7

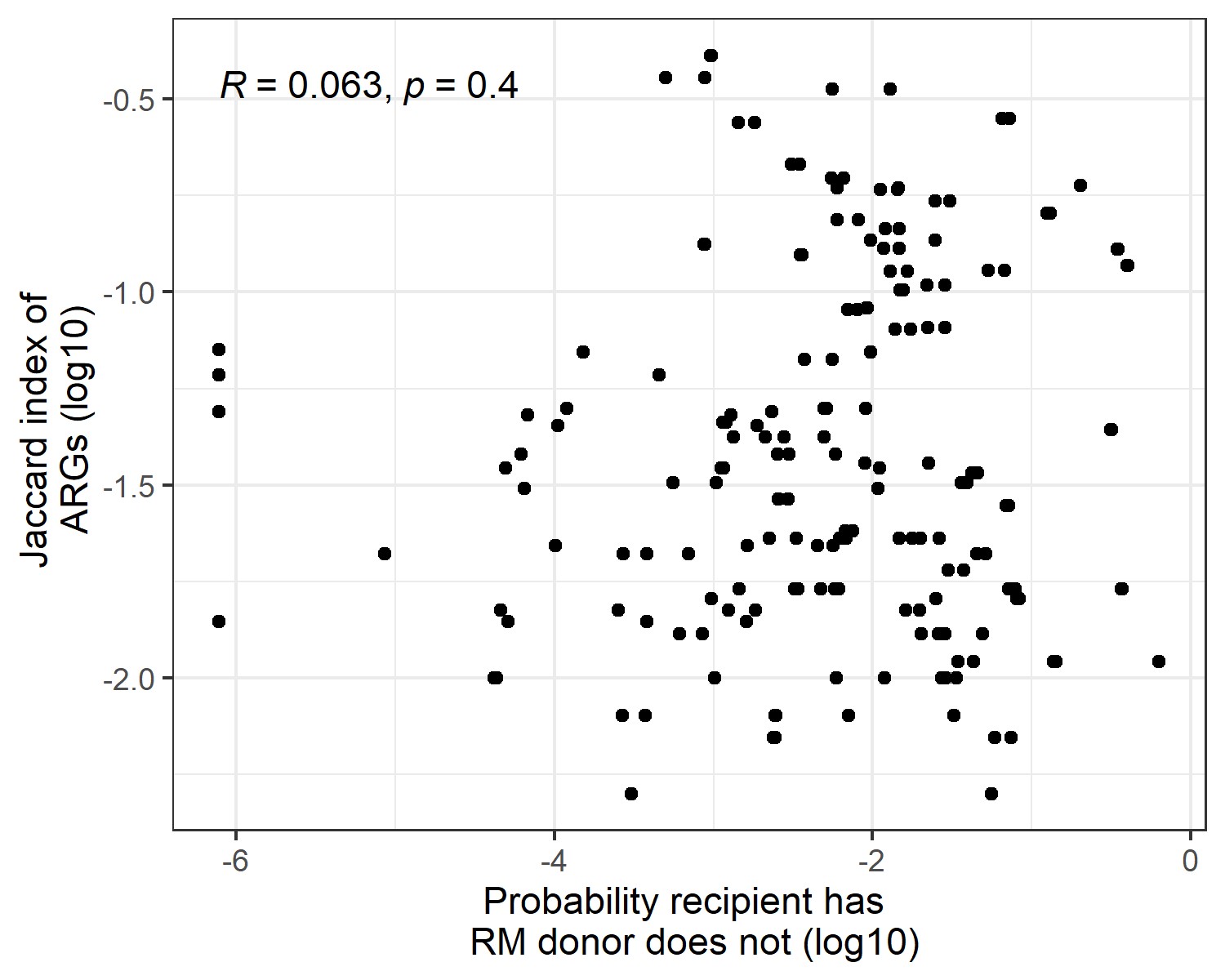
